# Large-scale neural recordings with single-cell resolution in human cortex using high-density Neuropixels probes

**DOI:** 10.1101/2021.06.20.449152

**Authors:** Angelique C. Paulk, Yoav Kfir, Arjun Khanna, Martina Mustroph, Eric M. Trautmann, Dan J. Soper, Sergey D. Stavisky, Marleen Welkenhuysen, Barundeb Dutta, Krishna V. Shenoy, Leigh R. Hochberg, R. Mark Richardson, Ziv M. Williams, Sydney S. Cash

## Abstract

Recent advances in multi-electrode array technology have made it possible to monitor large neuronal ensembles at cellular resolution. In humans, however, current approaches either restrict recordings to only a few neurons per penetrating electrode or combine the signals of thousands of neurons in local field potential (LFP) recordings. Here, we describe a new probe variant and set of techniques which enable simultaneous recording from over 200 well-isolated cortical single units in human participants during intraoperative neurosurgical procedures using silicon Neuropixels probes. We characterized a diversity of extracellular waveforms with eight separable single unit classes, with differing firing rates, positions along the length of the linear electrode array, spatial spread of the waveform, and modulation by LFP events such as inter-ictal discharges and burst suppression. While some additional challenges remain in creating a turn-key recording system, high-density silicon arrays provide a path for studying human-specific cognitive processes and their dysfunction at unprecedented spatiotemporal resolution.

---

Major technological advances in the past decade have led to a revolution in the neurosciences. Many research programs now routinely rely on the analysis of single-neuron action potentials from hundreds and even thousands of neurons, which provide a rich understanding of the coordinated activity of large neuronal ensembles that underlie sensory, motor, and cognitive operations ^1–4^. While these developments have been most pronounced in animal models, there have been parallel, albeit slower, advances in the ability to record from single neurons in humans. Single-unit recordings in humans have been performed since the mid-1950s ^5–8^, and were foundational in understanding the role of neural circuits in neurologic disease. For example, such techniques helped to establish an understanding of the relationship between basal ganglia dysfunction and Parkinson’s disease ^9^. There are currently at least four high-resolution neuronal recording technologies that can be used in human participants in acute, subacute, and even chronic settings. These include microwire bundles ^10, 11^, laminar microelectrode arrays ^12, 13^, microelectrode contacts arranged on a grid for use above the pia or on the shaft of a depth electrode ^14–16^, and the ‘Utah’ planar array of penetrating microelectrodes ^17–19^. Recent technological developments to move beyond more rigid structures include thin-film based systems with organic polymer electrodes (often referred to as micro-electrocorticography, ‘µECoG’), which provide access to surface recordings in the cortex ^20–25^ and even from the surface of the hippocampus intraoperatively ^26^.

While each of these approaches is nominally capable of recording action potentials from individual neurons, all are limited to capturing only 10-150 separable units (and typically well below 150) per device. In addition, these approaches rarely allow for high-quality isolation of action potentials from more than one neuron on a single recording channel (and vice-versa, rarely observe a single neuron from multiple channels). This constraint limits the quality of spike sorting and biases observations towards neurons with large and distinct waveforms with relatively higher spike rates. Improvements to existing recording systems should both increase the quantity of neurons an experimenter can record while simultaneously allowing for high quality spike sorting.

Demonstrations of neural recording systems using animal models have advanced at a substantially more rapid pace. A recent landmark for these advances has been the introduction of the silicon Neuropixels probe, a fully-integrated linear silicon microelectrode array with a single 10 mm shank (**Fig. 1a**), which is covered with microelectrode contacts at a 20 µm site-to-site spacing. The Neuropixels 1.0 probes can simultaneously record from 384 user-selectable channels distributed along a 24µm x 70µm x 10 mm shank. This system, introduced in 2017, has already enjoyed widespread adoption for recording in rodents ^1, 2^ and non-human primates ^3^, with continuing improvements to reduce the size and provide additional form factors ^4^.

**Figure 1:**
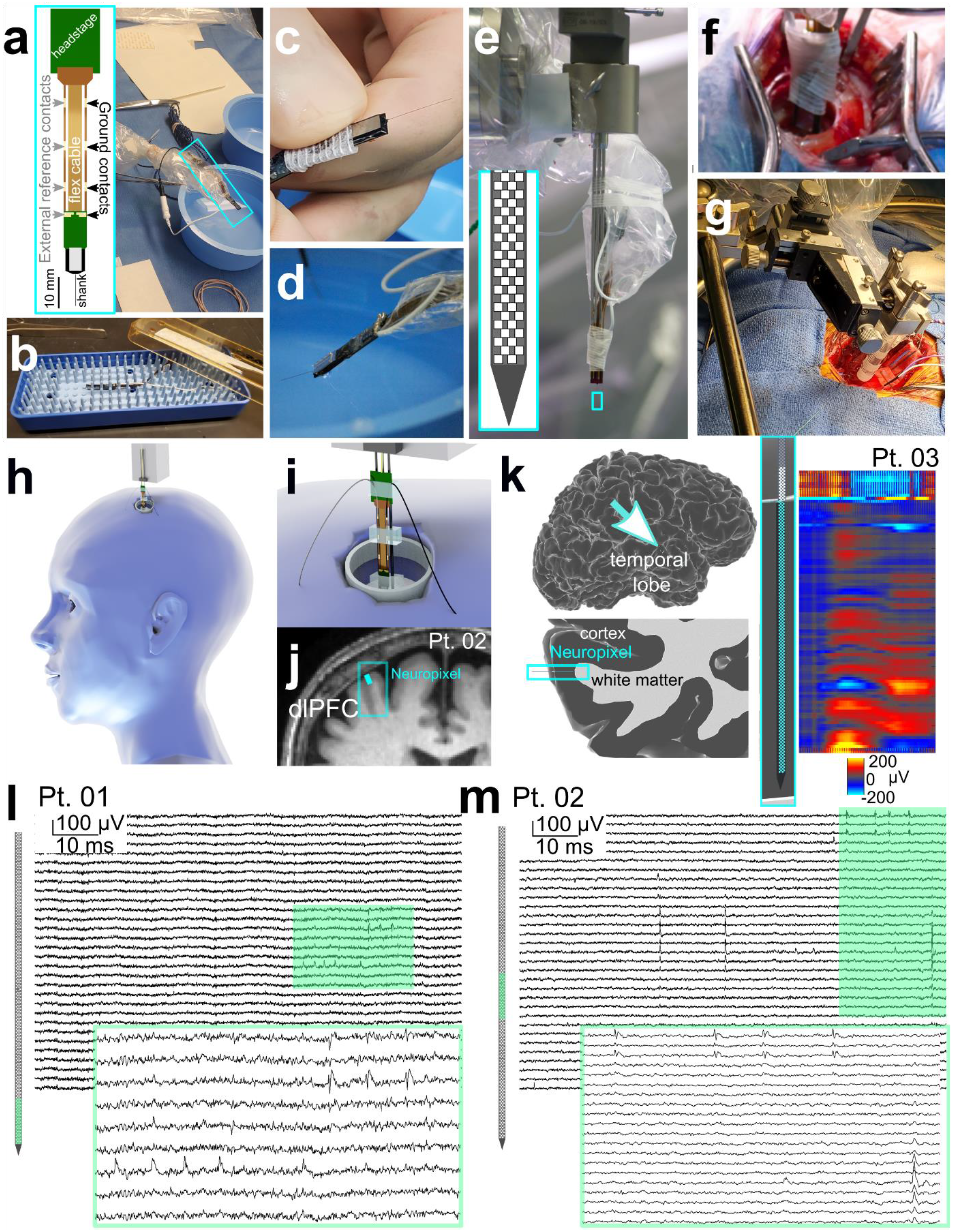
Human neocortical neurons recorded using Neuropixels 1.0-S probes. **a-d**: Diagram of the Neuropixels 1.0-S probe with a headstage and the ground and reference pads indicated (outlined in cyan, left) and preparation in a sterile field, with the probe outlined in cyan (**a**), set-up before electrode insertion, including opening the sterile electrode in the packaging, (**b**), handling and connecting to wires and visual inspection (**c**), and testing in saline (**d**). (**e**) Electrode attached to 3 sterile stylets on the ROSA robot for insertion. (**f**). Electrode inserted into the dorsolateral prefrontal cortex (dlPFC) through a burr hole using the ROSA robot. (**g).** Electrode inserted into lateral temporal cortex using a 3-axis micromanipulator attached to a Greenberg retractor. **h.** Three-dimensional model of the DBS burr hole location with a model of the Neuropixels probe. **i**. location zoomed in view on the three-dimensional view with the grey wire the reference and the black wire the ground. **j**. with the putative Neuropixels location overlaid on the preoperative MRI (top) during one DBS case which was mapped based on the implanted location of the DBS electrode (and the burr hole) and the angles of the Neuropixels probe based on the dimensions of the holder and burr hole as well as the closest visible cortical gyrus. **k**. Putative location and likely depth of the electrode in an open craniotomy case for epilepsy surgery in the lateral temporal cortex (left two columns), with the depth informed by the electrophysiology, where the LFP shows a clear difference between superficial electrodes and deeper contacts, as highlighted here in a color scale indicating voltage. **l-m**. Example recording from participant 1 (Pt. 01) and participant 2 (Pt. 02) in the dlPFC across multiple channels, with action potentials shown extending across multiple channels. The light green filled in box in the background traces are then expanded in the green-outlined voltage traces in the foreground. In **a, e**, **j**, and **k**: cyan rectangles are highlighting the location of the Neuropixels probe.

Recording single unit neural activity in humans is increasingly common in both research and in clinical care, including the use of real-time neurophysiology to guide deep brain stimulating electrodes for neurological disorders like Parkinson’s disease ^9, 27–34^. Research to develop clinical brain computer interfaces for individuals with severe speech and motor disorders has used Utah arrays to record small populations of single neurons and groups of several neurons. Utah arrays have proven instrumental in enabling basic neuroscience and clinical studies at the level of single neurons (though not necessarily well isolated single units) and at the population level. These studies range in application from decoding handwriting and controlling prosthetic limbs and reanimating upper extremities to providing sensory feedback (via intracortical microstimulation) ^35–44^. Adapting the Neuropixels recording system for use in clinical research presents a viable approach for building on these prior demonstrations of the utility of cellular-resolution neural recordings for clinical and neuroscientific applications.

We sought to develop a suite of techniques for using Neuropixels probes to record brain activity acutely during clinically-indicated neurosurgery. Scaling up the size and quality of neural population recordings is a crucial prerequisite step to enable novel fundamental neurophysiological and clinical investigations. For example, detailing the cellular-scale mechanisms underlying epilepsy ^27, 30^ or understanding changes in cellular interactions induced by the presence of tumor cells ^45, 46^ or as part of an advance brain computer interface would be drastically improved via high-density recordings. The versatility of the Neuropixels probe allowed us to record activity both during the placement of deep brain stimulators (DBS) and during open craniotomies for removal of brain tissue for the treatment of epilepsy and brain tumors.

Here, we demonstrate a set of methods, including a new, thicker variant of the Neuropixels probe, which enables acute experiments in human patients. We describe solutions to challenges in sterilizing probes, electrical isolation in the operating room, and brain pulsation after craniotomy. We used this approach to record from populations of neurons in dorsolateral prefrontal cortex and temporal lobe, and to subsequently characterize the electrophysiological diversity of neurons in these regions of human cortex. We observed a wide diversity of extracellular waveforms (assigned into different putative cell type groupings, validated using three separate clustering techniques). In contrast with prior observations ^47^, firing rates only differed slightly between narrow- and wide-waveform amplitude units. We also found single cell activity covaried significantly with epileptiform activity as well as anesthesia-induced burst suppression. Taken together, this new version of the Neuropixels probe and a set of methods to use them in interoperative settings lays the foundation for expanding our understanding of human cortical function in both health and pathology and in developing neurorestorative technologies.

## RESULTS

Here, we report successful recordings from the cortex of temporal and frontal lobes in patients undergoing brain tissue resection to treat epilepsy (N=1, under general anesthesia, lateral temporal lobe) or during the implantation of DBS leads to treat movement disorders (N=2, one awake and one under general anesthesia, dorsolateral prefrontal cortex) using Neuropixels probes. We also report unsuccessful recordings – and lessons learned -- from six cases performed while developing these approaches (**Supplemental Table 1; Supplemental Figure 1;** see **Methods**). Unsuccessful recordings were either due to electrode fracture (N=2, with the devices and pieces fully recovered; **Supplemental Table 1**) or excessive noise during the recordings (N=4). Two different types of Neuropixels arrays were tested: a thinner Neuropixels 1.0 probe (thickness: 25µm, width: 70µm, length: 10 mm) was utilized in two of the six failed attempts; neither of these yielded stable recordings. Given the challenges introduced by using such a thin probe in the operating room context, we developed a variation of the Neuropixels 1.0 probe, featuring a 100µm thick shank (Neuropixels 1.0-ST: thickness: 100µm width: 70 µm, length: 10 mm). This version enabled considerably easier insertions and robust use during neurosurgical cases, easing the process of inserting the probe and reducing the risk of mechanical failure. This probe version, combined with an improved grounding and reference electrode configuration, enabled us to observe spiking activity from populations of isolatable single neurons in three participants (N=3, **Supplemental Table 1; Supplemental Figure 1-2;** see **Methods**).

Since the Neuropixels probe is typically used for small animal neurophysiology, five technical developments were needed to translate this device to intraoperative use in people. These included 1) sterilization with ethylene oxide and maintaining sterile conditions in the operating room, 2) mounting the probe to a neurosurgical robot or a sterile micromanipulator, and 3) stereotactically guided insertion through a burr hole or craniotomy window. These techniques, some of which were informed by our previous experience adapting Neuropixels to NHPs ^3^, are described in further details in the **Online Methods**. A fourth, and crucial, consideration was the identification and reduction of sources of noise in our recordings in the operating room (OR), which are considerably larger and harder to control than in experimental laboratory settings for animal research. We performed tests both during the neurosurgical cases as well as in the OR without a patient to identify the external sources of noise (e.g. anesthesia IV pumps) as well as internal sources of noise (e.g. we found that in this environment ground and reference should be separate and not tied together has is often required in mouse studies; see **Methods; Supplemental Fig. 1**). Fifth, as it was not possible to suppress the brain due to patient safety considerations and shear stress on the Neuropixels probe (see **Methods**), a semi-automated post-hoc registration method was developed to allow for single units to be stably isolated and tracked over time.

Single unit waveforms were observed both on individual Neuropixels probe channels and across multiple channels (**Fig. 1-2**). However, we also observed considerable modulation of the voltage recording related to motion of the brain. This motion primarily results from respiratory and cardiac rhythms, which cause the surface of the brain to move relative to the probe. We observed these movement-related changes in both the LFP and action-potential bands (**Fig. 2a-d**). To confirm that the movement present in the neural recordings was due to tissue movement, we matched the neural recording itself, specifically the LFP band, to the synchronized audio of the EKG and video of the brain (**Fig. 2b**).

**Figure 2:**
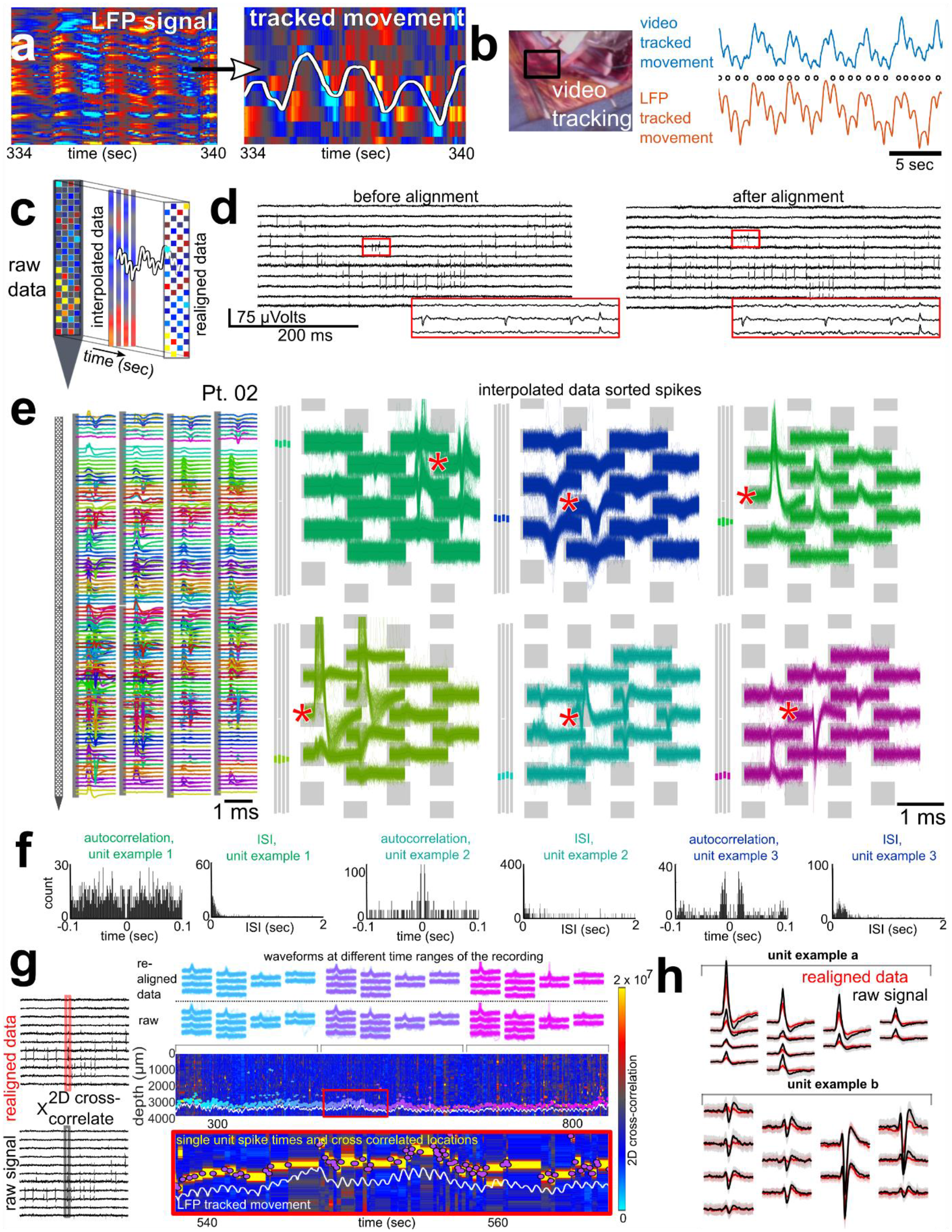
A variety of waveform types and shapes recorded in human neocortex with Neuropixel probes. **a.** Illustration of evidence of tissue movement relative to the electrode recordings in the LFP (shown in red-blue color scale with the range in µvolts shown in **c**). This is quantified by manually tracing these “band shifts” using the Blender program, followed by detection of these movements in the LFP and tracking of these movements across channels (white line, second to rightmost plot). **b.** Validation of movement being reflected in the LFP channel shifts. Left: video of the intraoperative recording and the pumping evident in the CSF surrounding the electrode was tracked through time. Right: Simultaneous traces of the video tracked movements and the LFP-tracked movements in the same patient (Pt. 03) at two different scales. Generally, the magnitude of the movement artifact was on the order of 80-100 µm. **c.** Series of steps taken for adjusting for movement and interpolating the data. **d.** High frequency (action potential) frequency signal before (left) and after (right) adjusting for movement effects. **e.** Example unit waveforms (each color is a separate unit, * indicates the largest waveform per unit and this was used for further waveform measurements). On the left, original waveforms are overlaid relative to the recorded channels, with the grey bars to the left indicating the location of the units along the probe. **f**. Example autocorrelograms and inter-spike-interval (ISI) distributions for three different units. **g**. Approach for cross correlating the movement-corrected neural data and the raw signals by cross correlating the 12 channels of the sorted spikes with the raw data (left). Top: tracked single unit across 12 channels identified in the realigned data (top row) as well as the raw data (second row; following cross correlation), with waveforms from single spike times overlaid from different epochs of time (indicated by the brackets beneath the waveforms). Cyan traces: first third of the recording, purple traces: middle third of the recording; magenta traces: last third of the recording. Bottom: Cross-correlation values (shown in color map) which was paired with the LFP tracked movement (white lines) allowed tracking of individual units along the raw channel data (bottom, right, purple dots). **h**. We could then correlate the 12 channels with the waveforms on a per-unit and per channel level and overlay the unit waveforms for the raw data (black lines) and over the average units sorted from the realigned data (red lines).

Critically, the high spatial density of Neuropixels electrodes allows for post-hoc motion correction, and high-quality spike sorting. Estimating the timepoint-by-timepoint brain position relative to the probe was best achieved using the LFP channels. (**Fig. 2a-c**; **Supplemental Fig. 3**). We compared multiple approaches for correcting tissue motion and determined that the optimal approach was to use semi-automated tracking (See **Supplementary Methods**) taking advantage of the high-density sampling and the high temporal resolution of the LFP (**Supplemental Fig. 3-4**). The motion contained multiple dynamics with one peak around 1.5 Hz which shifted with time (possibly with heartrate), a variable slow oscillation (∼0.15 Hz) that dampened over time (possibly as the electrode settles into the tissue), and slow drift (∼0.01-0.02 Hz) that has different timing dynamics as shown with a spectrogram of the detected motion (**Supplemental Fig. 4**). This combination of frequencies proved challenging for fully automated approaches that rely on a >1-second time scale and spikes for motion correction ^48^ and were not compensated for by the drift correction offered by Kilosort 3.0 ^4, 49^ (**Supplemental Fig. 3**).

Following this movement alignment, we sorted waveforms into clusters using Kilosort 3.0 software ^4, 49^ and extracted an average of 201 ± 151.04 clusters from the recordings. Each cluster presumably represents action potentials generated from a single neuron or very small number of neurons and has both temporal characteristics (waveform shape) and spatial characteristics (the pattern of waveforms across multiple recording channels). Data from the two participants undergoing DBS implants contained similar numbers of clusters (with the Neuropixels recording from the dorsolateral prefrontal cortex, Pt. 01, not awake: n= 262; Pt. 02, awake: n= 312), while the third participant undergoing an anterior temporal lobectomy yielded considerable fewer (Pt. 03, not awake: n= 29). The waveform clusters were then classified as either multi-unit activity clusters (MUA; Pt. 01: n= 60; Pt. 02: n= 134; Pt. 03: n = 10) or single units as described in detail below (Pt. 01: n= 202; Pt. 02: n= 178; Pt. 03: n = 19; **Supplemental Fig. 5-6; Supplemental Table 1**).

As the single units could be captured both vertically and laterally on a fine spatial scale, the advantage of the contact-to-contact 20µm distance relative to the known sizes of human cortical neurons (approximately 20 µm at the cell body; **Fig. 2b, 2; 3a-c; Supplemental Fig. 5**) combined with the checkerboard arrangement of the Neuropixels array allowed a sub-sampling of the extracellular single unit action potential. As such, we could reliably trace the location and identity of single cells across the recording, an approach which would be limited if we could only access a single channel at the >150 µm resolution of other single neuron recording devices (**Fig. 2d; 3a-c;** ^12, 17, 27, 50^). Specifically, we found an average of 52% of units had significant (5 STD above baseline voltage) deflections across multiple channels (Pt. 01: 90 units out of 202 units; Pt. 02: 103 units out of 178 units; Pt. 03: 10 units out of 19 units; Fig. 2; Supplemental Video 1). We could then take advantage of this feature to determine if the movement correction and interpolation could influence the detection and tracking of single units and whether the same identifiable single unit waveforms could be found in the raw data throughout the recording. We performed a two-dimensional cross correlation between the sorted waveforms and the raw action potential (high frequency band) data (**Supplemental Fig. 4**). The process involved cross correlating the 12 channels of the sorted spikes from the movement-corrected data stepwise channel set by channel set through the raw data (**Supplemental Fig. 4**). We found that we could track the spatial location of individually identifiable units (not simply the largest waveforms) through time and spatially in the raw data along the electrode, we were able to calculate the average movement found per patient (**Fig. 2g; Supplemental Fig. 4**). After these steps, we found high correlations on the per-waveform and channel level across units and participants (mean Pearson’s correlation per waveform per channel; Pt. 01: 0.64±0.197, maximum: 0.99; Pt. 02: 0.65±0.254, maximum: 0.99; Pt. 03: 0.50±0.125, maximum: 0.60; **Fig. 2h**). Overall, the correlation between the raw data and the sorted spikes increased with increasing number of spikes (Pearson’s correlation between number of spikes and correlation values: Pt. 01: rho=0.27; p=0.0001; Pt. 02: rho=0.33; p<0.0001; Pt. 03: rho=-0.35; p=0.137; Supplemental Fig. 4).

Finally, we found that most units could be detected throughout the recording in the raw data (mean range of time the units are detected: Pt. 01: 480.1±129.70 sec; Pt. 02: 438.8±94.92 sec; Pt. 03: 517.9±165.90 sec; proportion of the recorded time the units were detected: Pt. 01: 0.82±0.221; Pt. 02: 0.77±0.166; Pt. 03: 0.71±0.229; **Fig. 2g; Supplemental Fig. 4; Supplemental Video 1**). Both with the cross correlated tracking of the single units and the LFP movement detection, we found the movement variance was higher for the awake case over the two not-awake cases (LFP variance in movement: Pt. 1: 2.31; Pt. 2-awake case: 26.15; Pt. 3: 4.47; Cross-correlated unit maps: significant differences between patients; Kruskal-Wallis multiple comparisons test; Chi-sq.=399.39; p<0.00001; **Fig. 2i; Supplemental Fig. 4**). Last, we found that the mean velocity of the movement as calculated from the LFP was 76.5±27.30 µm/sec (**Supplemental Fig. 4**).

After confirming that identifiable single units could be tracked both in the movement-corrected and raw data sets, we observed a diverse set of spike waveforms within and across participants. Broadly speaking, the waveform clusters could initially be subdivided into those with a dominant positively deflecting or negatively deflecting peaks as well as double-peaks and triple-peaked waveforms. A neuron’s waveform changes depending on the location of the recording electrode with the shape of the waveform not an invariant property of the neuron, but a function of the both the neuron and the recording geometry ^51, 52^. As would be expected from electrically observing a neuron from multiple vantages ^51^, some isolated units had more than one type of waveform across the different channels. Enabled by the ability to record simultaneously from multiple closely-spaced electrodes, a single putative unit’s voltage changes during action potential generation could be detected as differently shaped waveforms. For example, there were units with positive peaks on some channels and, simultaneously, negative peaks on other channels or double peaks on one channel and single peaks on other channels (**Supplemental Fig. 5**). Finally, different units were either highly localized to 2-3 channels while others were widely dispersed across electrodes, with waveforms of various durations and amplitudes (**Fig. 1e-f; Fig. 2c,g; Supplemental Fig. 5**).

With the ability to record simultaneously from multiple closely-spaced electrodes, unlike with other state of the art microelectrode recording devices (**Fig. 3a-b**), we could spatially sub-sample and confirm that the voltage generated per single cell during the putative action potential was detected as differently shaped waveforms (**Fig. 2e-f; Supplemental Fig. 5; Supplemental Video 1**). These phenomena have been previously observed in non-human primates (NHPs; ^49^ and rodents ^1, 2, 4^), but such a high-resolution view of electrophysiological waveform diversity has not been previously available *in vivo* in humans, particularly since the smallest contact-to-contact spacing regularly used in human studies is 150 µm; a spacing that is considerably larger than a single neuronal cell body and the 20 µm contact to contact spacing in the Neuropixels probe (**Fig. 3a-b**). In the following sections, we describe in more detail the classifications of different unit types of the spike sorted units (only using Kilosort 3.0), followed by resulting electrophysiological findings of the firing and spatial properties of these different classified groups of units, preliminary neurons’ spiking relationships between each other in the recording, and spiking relative to events found in the LFP, including epileptiform activity and anesthesia-induced burst suppression.

**Figure 3:**
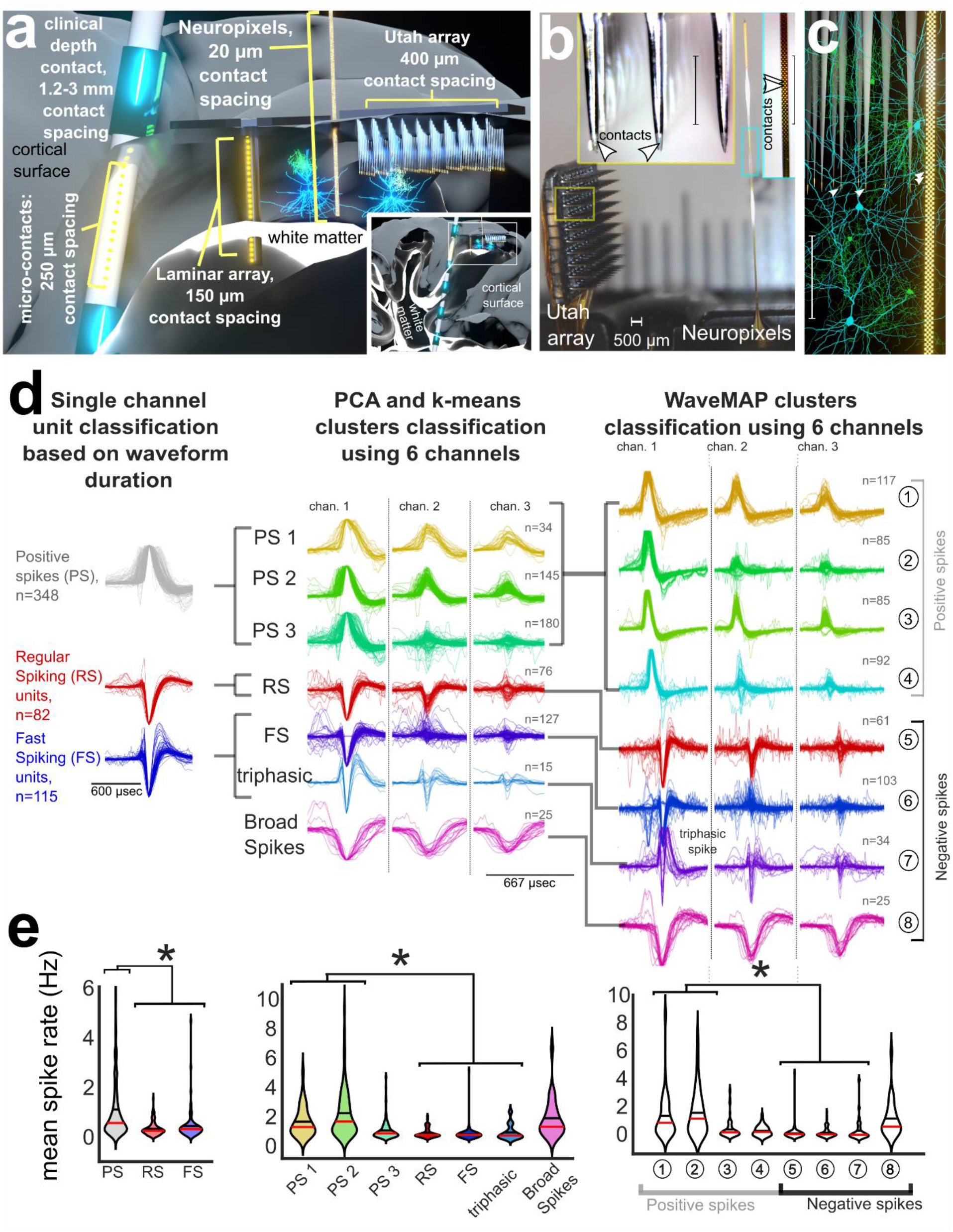
Classifying waveforms based on spatial and temporal features. **a**. Illustration of different types of state-of-the-art intracranial microelectrodes implanted in cortex, with a zoomed out view of the clinical depth lead and the electrodes in cortex in the inset box in the lower right corner. **b**. Photograph of the Utah array side by side with the Neuropixels array with the yellow (Utah array) and cyan (Neuropixels) insets at the same scale. **c**. Reconstructions of human pyramidal cells and inhibitory interneurons with cells from NeuroMorpho.Org ^61–65^ relative to the Neuropixels array and 1.5 mm Utah array. **b-c**: Scale bar 500 µm. **d**. Left: average waveforms of each unit using the single channel classifications. Middle: average three channels per unit (each column is a channel), multichannel classification based on the principal components and k-means clustering across patients. Right: average three channels per unit (each column is a channel), multichannel classification based on the WaveMAP approach across patients.

### Cell type classification based on extracellular waveforms

We observed a diversity of waveforms. While there could be a stochastic element in that the waveform direction and shape could be more dependent on the location of the electrode relative to the cell body ^51^, there are a growing set of studies arguing that positive deflecting waveforms represent axonal action potentials as shown in Neuropixels recordings in other species ^2, 53, 54^. In addition, we wanted to link the current Neuropixels spatially subsampled action potentials to past human single unit studies which could only use single channels at lower contact-to-contact spatial resolution to record action potentials (>150 µm contact to contact spacing; ^17, 27, 50^). We decided to categorize the cluster waveforms taking into account each of these points using a three-pronged approach, particularly to emphasize the strength of the Neuropixels device in sampling multiple putative cell types or waveform morphologies.

We used multiple classification techniques, including both supervised and unsupervised algorithms, to determine if these methods can reliably identify distinct extracellular waveform classes from the sorted units (which could relate to different cell types) across patients and across classification schemes (**Fig. 3d**, and **Online Methods; Supplemental Fig. 6-9**). First, we used a single-channel approach routinely employed by researchers to classify putative inhibitory and excitatory neurons. This single-channel classification was based on the polarity and half-peak waveform width of the largest waveform for each unit (i.e., from the one channel with the largest waveform for that unit) using published unit type definitions as used in most human single unit studies to date (**Fig. 3d**, and **Online Methods; Supplemental Fig. 7-8** ^49, 50^). This classification produced broad classes of units but with considerable variation within each class such that we did not find significant differences in unit features for a number of metrics.

Next, to take advantage of the Neuropixels’ high-density spatial sampling of action potentials, we performed principal component analysis (PCA) followed by k-means clustering to identify distinct classes of units with stereotyped waveform shapes, taking into account 6 channels per unit instead of just the one (**Fig. 3d; Supplemental Figure 6**). Finally, though there is no ground truth available, we repeated this analysis using an alternate unsupervised, classification algorithm called WaveMAP. The WaveMAP algorithm automatically identifies and separates cell types based on waveform shapes across six (or 12) channels utilizing Universal Manifold Approximation and Projection (UMAP) dimensionality reduction combined with Louvain clustering^55^. For each of these approaches, we identified differences between the putative unit types in their waveform characteristics as well as their spatial location along the probe shank.

Each classification improved the separability of the classes of units, with the different classes showing common features between single channel, PCA, and WaveMAP approaches (**Fig. 3d; Supplemental Fig. 9**). For instance, all three approaches produced the same sets of classifications of negatively deflecting regular spiking (RS) and fast spiking (FS) units as well as similar classifications of positively deflecting waveforms between the PCA/k-means approach and WaveMAP.

### Single channel classification of waveforms based on duration and peak direction

In accordance with common practice, we used single-channel metrics (waveform half-peak width and duration (valley-peak) to separate units into three classes: positive spikes (PS; n=348), versus negative spikes with the later subdivided into negative fast spiking (FS; n=115) or negative regular spiking (RS; n=82) single units based on the peak-trough duration of the largest waveform across channels per unit and the polarity (asterisks in **Fig. 2e;** see **Online Methods**) ^47, 50, 54, 56–58^. As positive deflecting waveforms have been hypothesized to be the results of axonal action potentials, we separated positive and negative spikes ^2, 53–55^. We also included a fourth category of multi-unit activity (MUA) where the waveforms were mixed across the single unit cluster and the autocorrelation and inter-spike-interval (ISI) distributions were more representative of MUA, which were more uniform in distributions as opposed to single unit distributions (**Fig. 3d**). The peak-trough duration cutoff for FS versus RS units was at 300 µs, which is similar to the cutoff reported previously ^47, 59, 60^. This boundary was also near a boundary in a presumed bimodal distribution of peak-trough durations for events with negative polarity (purple line, **Supplemental Fig. 7**).

Each putative class of neuron displayed differences in waveform features and spike rates (**Fig. 3d, Supplemental Fig. 7-8**). First, PS units generally had wider waveforms compared to the RS and FS units. The half-peak width was 167.3±55.4 µs for PS and 133.8±70.9 µs for FS + RS, while spike duration was 494.2±175.1 µs for PS and 349.6±222.2 µs for FS and RS; **Fig. 3d; Supplemental Fig. 7)**. These unit types also showed significant differences in repolarization slope and recovery slope as well as the peak-trough ratio (**Supplemental Fig. 7**). Between these three categories of units (PS, RS, and FS), the PS unit mean firing rates were significantly higher than those of RS and FS units (p<0.001; Wilcoxon rank-sum test; RS: 0.3± 0.27; FS: 0.4±0.59; PS: 1.1±1.35 Hz mean firing rate). FS units trended, though non-significantly, towards slightly higher firing rates than RS units (p>0.05; Kruskal-Wallis multiple comparisons test; **Fig. 3e**). Spike rates were higher in the awake case overall. However, there was no significant difference in spike rates between the FS (narrow waveform) and RS (wide waveform) units in either the anesthetized cases (p=0.0705; Wilcoxon rank sum test) or the awake cases (p=0.0901; Wilcoxon rank sum test). The peak-trough ratio was significantly lower for the FS and RS units than PS units (PS: 4.8±2.0 RS: 3.9±1.7; FS: 4.1±1.8; p<0.001; Kruskal-Wallis multiple comparison test). For the other waveform measures, the recovery slope was higher in absolute amplitude for the RS units than the FS and PS units, while the repolarization slope was larger for the PS units than RS or FS (**Supplemental Fig. 7**). These differences in waveform characteristics resemble what has been reported in previously in other species (mice, cats, and macaques)^2, 47, 54^. However, this single channel classification explicitly ignores the spatiotemporal features of extracellular waveforms afforded by Neuropixels recordings high resolution sampling (**Fig. 3a-c**).

### Multichannel classification of units using PCA and k-means clustering

For multi-channel classification we also used an unbiased clustering approach (**Fig. 3d**) by applying principal components analysis (PCA) to the mean waveforms for each unit (using a larger feature vector that includes data from multiple electrode sites) per class, followed by k-means clustering (**Fig. 3d; Supplemental Fig. 6**). This analysis identified seven classes of units using the waveforms from the first six channels of each unit (with channels reordered-from largest to smallest waveform per unit; **Fig. 3d**). The resultant classes further subdivided the categories identified using single channel categorization varying temporal and spatial distributions as well as triphasic waveforms, with two of the classes closely resembling the RS and FS designations from the previous analysis, presented above (**Fig. 3d**). This alternative classification schema also revealed there were increased firing rates for two of the positive spikes (PS1-2) relative to FS and RS units. In addition, FS firing rates trended higher than RS firing rates (though this did not reach statistical significance). We also found similar trends and differences between the PS and other waveforms with increased PS peak-trough ratios and varying repolarization and recovery slopes per waveform classification (**Supplemental Fig. 7**).

### Multichannel classification of units using WaveMap

Finally, we utilized an automated, non-linear method for classifying extracellular waveforms, with the goal of identifying additional classes based on smaller and more nuanced features of the waveforms ^55^. Using the WaveMAP algorithm^55^, we found that specific waveform types were identifiable across participants. We further subdivided the units into data from each participant. Even with this subdivision, after performing WaveMAP on each data set, the same WaveMAP classifications could be found with these independent classifications per participant. Negative RS and FS-like units were present in all three patient’s data, while the positive large waveforms (PS) appeared in two of the three cases (**Fig. 3d; Supplemental Fig. 9**). We therefore pooled the waveforms across patients and classified waveforms from across the six channels with the largest amplitude waveforms of each unit using WaveMAP which revealed eight total classes, representing four positive spike (PS) classes and four mostly negatively deflecting neuron classes. The four negatively deflecting spike classes included FS-like, RS-like, triphasic, and broad classes (**Fig. 3d**). As with PCA and k-means classification, we found little difference in the number of classes when we included either six or 12 channels with the largest amplitude waveforms in the classification. In addition, as we found with the other neuron classification approaches, we found similar differences in spike rates for different PS classes (including two of the three PS classes having higher spike rates than the RS, FS, and triphasic neurons) as well as the RS and FS-like neurons including subsets of PS neurons (corresponding to a unique waveform class) exhibiting higher spike rates (**Fig. 3e**).

### Differences in spatial spread of single channel and multichannel waveform classes

An important feature of the Neuropixels probe is the ability to obtain high local spatial sampling, which permits tracking of individual spike propagation and has been used to argue for extracellular identification of back propagating action potentials (**Fig. 3a-c**)^1, 2, 4^. Using published metrics and code ^2^, we calculated the spatial spread and velocity of the waveforms among the three identified categories of waveforms (**Supplemental Fig. 8**). In the single channel classification, we found the spatial spread for the FS and RS units was significantly lower than for the PS units (p<0.001; Kruskal-Wallis multiple comparison test). This trend held when looking at the multichannel classification (PCA/k-means and WaveMAP) with some PS units with higher spatial spread than RS, FS, and triphasic units (**Supplemental Fig. 8**).

Finally, mapping the different waveform classes to their location along the probe relative to the surface of the cortex, we found PS units were observed at sites throughout the shank of the probe (and throughout the cortical layers) while the RS and FS type units appeared to be concentrated deeper in the cortex. This spatial distribution of these broad cell categories likely reflects the inhomogeneous distribution of cell types and action potential propagation patterns throughout cortical lamina, though a precise mapping of PS, RS, and FS waveform groups to more detailed cell classes is impossible to perform using extracellularly recorded waveforms alone (**Supplemental Fig. 8**). That said, we found more subtle differences among the PS groups as well as the RS, FS, and triphasic spike groups with the multichannel classifications (PCA/k-means and WaveMAP) in the unit depth (along the Neuropixels probe) for different PS units as well as the RS and FS-like units. A subset of PS units were located in more superficial cortical lamina relative to the negative spikes and even other PS units (**Supplemental Fig. 8**). However, in none of the waveform classes (single or multichannel) were there a significant difference in the velocity of the waves above or below the peak waveform. This pattern suggests that the action potentials being recorded here are localized and possibly generated at the soma (**Supplemental Fig. 8**) ^2^.

In sum, we found that there was strong agreement in the three different unit classifications between the single channel and multi-channel classification approaches, though the multichannel classifications (PCA/k-means and WaveMAP) revealed more putative cell subtypes among the units which were supported by significantly different firing rates, spatial distribution, depth, and waveform features.

### Timing and interactions among single units

We could observe units within seconds of insertion on in the recording, though there was generally a ∼2 minute period of time when both single unit and MUA spike rates would slowly increase in all three participants’ recordings before levelling off at a seemingly consistent rate (**Supplemental Fig. 10**). We mapped the WaveMAP classified single units as well as the MUA classes to the electrode depth and the duration of the recordings with the different WaveMAP classes observable throughout the recording and across the Neuropixels shank (**Supplemental Fig. 10**).

To determine, as a proof of concept, whether there were interactions between units through time, we performed pairwise covariance analyses between the spike times of different pairs of units and found correlations between individual units in two data sets (Pt. 01 and Pt. 02; **Supplemental Fig. 10**) ^66, 67^, suggestive of inter-unit circuit activity. We found that some individual units covaried with each other in time (within a 0.1 sec time window), even between different classes (**Supplemental Fig. 10**). Across the WaveMAP classes, we found an average of 6.9 ± 2.58% of all pairs had a significant covarying relationship (at least one binned peak in the covariance calculations 8 STD above the baseline covariance levels across the recording). The average absolute lag between pairs with significant covariances was 13.9 ± 2.0 ms (Pt. 01: 14.1 ± 14.2 sec; Pt. 02: 15.4±14.9 sec; Pt. 03: 12.7± 13.6 msec). Of these comparisons, we found that 58% of the pairs with significant covariances showed absolute spike lags > 5 ms and 46% were even longer than 10 ms (Pt. 01: >5ms-60%, >10ms-46%; Pt. 02: >5ms-65%, >10ms-52%; Pt. 03: >5ms-49%, >10ms-42%). However, a challenge in interpreting these temporal lead/lag relationships is that epileptiform activity or general anesthesia induced burst suppression could have induced correlated class activity in 2 of the 3 participants (**Supplemental Fig. 10**). We anticipate more detailed pairwise relationships between units can be identified in future studies without these confounds ^68–70^ or using further analytical techniques accounting for the specific effects of anesthesia ^71–73^.

### Neuropixels recordings can reveal spike-field relationships

Numerous questions in clinical and cognitive neuroscience have been addressed by examining the relationships between LFPs and the timing of spikes ^74–76^, even in the short period of time (15 minutes) of these recordings using single channel lowered electrodes ^32, 77, 78^. To that end, we next sought to determine whether the single and multi-units detected using Neuropixels could also be used to extract spike-field relationships (**Fig. 4**). We first movement-corrected the Neuropixels LFP recordings using the same manual approach as for correcting spiking activity which revealed local field voltage reversals along with the Neuropixels depth (**Fig. 4a**).

**Figure 4.**
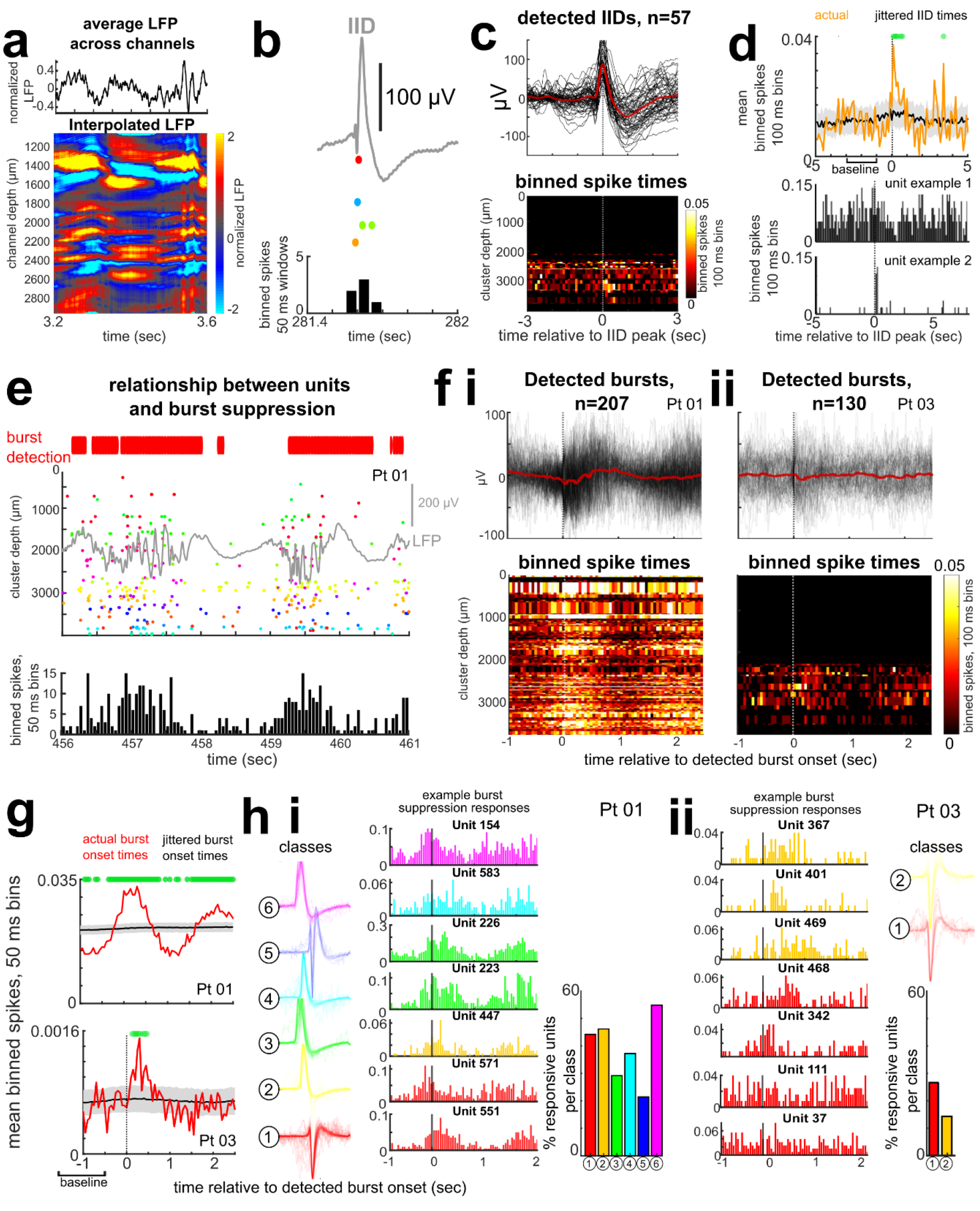
Relationship of units to the local field potential (LFP) events and epileptiform discharges. **a.** Example LFP averaged across the electrode (top) and along the depth of the Neuropixels probe after interpolating the LFP using manual registration. **b.** Inter-ictal epileptiform discharge example (IIDs, grey line) along with single unit activity (colored dots represent spike times for different units) through time in Pt. 03, and binned spike times (below). **c.** Top: Individual (black traces, n=57) and average IIDs (red line). Bottom: Binned spike rates of individual units along the Neuropixels depth (y-axis), with the brighter pixels indicating increased activity aligned to the peak of the IIDs. **d**. Individual unit example spike timing relative to the peak of the IIDs. **e.** LFP burst suppression in a participant under general anesthesia, as shown by the grey line (LFP) along with the detected single unit activity (raster plot in the middle figure; each color represents a different unit) and binned into 50 ms windows (bottom). Red bars on top of the figure: automated burst detection algorithm ^81^. **fi-ii.** Detected burst onsets relative to binned single unit activity along the Neuropixels probe in two different patients (Pt. 01, **i**, and Pt. 03, **ii**). **g.** Binned spike rates relative to the burst onset (red) compared to time points where the burst onset was jittered randomly around a normal distribution of time ranges (black line). Green dots: spike rates (mean across units) which are significantly different compared to the −1.5 to - 0.5 sec before burst onset. **hi**. Left: Units separated into 6 classes for one patient (Pt. 01), color-coded per class. Middle: example unit activity relative to the 1 second before and 2 seconds after the burst. Some units increase spiking before, during, or after the burst. Right: Bar graph indicating the percentage of units per class with significant changes in firing rate relative to the burst onset. **hii**. Units separated into 2 classes for one patient (upper right, Pt. 03), color-coded per class, and example unit activity with changes in spike rates relative to the 1 second before the burst onset, with some units increasing spiking before, during, or after the burst. Below right: bar graph indicating the percentage of units per class with significant changes in spike rate relative to the burst onset.

Even without interpolating and re-aligning the LFP, we could identify interictal epileptiform discharges (IIDs) in the recording from the lateral temporal lobe before tissue resection for epilepsy (**Fig. 4b**). After confirming the presence of the IIDs using both automatic approaches ^79^ and visual confirmation by a trained epileptologist (SSC), we then aligned the single unit and multiunit activity to the peak of the IIDs (n=57). Single units increased their firing rate around the peak of the IIDs, even in recordings with sparse firing (**Fig. 4c**). To verify this change, we jittered (centered on a normal distribution) the timing of the IID peak relative to the unit times and found that the increase in spike events around the actual IID peaks was significantly above the binned spike times around a jittered IID peak (**Fig. 4d**). Further, we compared the average binned spike counts (across trials per time step) to a baseline −2 to −0.5 sec before the IID and found a significant increase in spikes in the half second after the peak of the IIDs (p<0.005; Wilcoxon rank sum test). Some units increased at the IID onset or only fired during the IID peak, while other units decreased firing around or after the IID, though we did not find a correspondence between unit class and spike rate modulation around the IIDs. This varying pattern amongst units has also been reported in Utah array recordings ^13^.

In two cases, the participants were under general anesthesia, which produced a typical burst suppression pattern in the average LFP (**Fig. 4e-f**; **Supplemental Table 1**; ^80^). Single unit firing on the whole increased during the bursts and decreased during the suppression periods in the LFP. The firing rate from −1.5 to −0.5 seconds before the onset of a detected burst relative to the −0.5 to +1 seconds after the onset of the detected burst was significantly increased. The unit firing during the bursts was significantly higher compared to the burst-triggered unit firing calculated in a time-shuffled data set, where the burst time onset was shuffled in time relative to the spike data (1000 shuffled tests, **Fig. 4g**). This population change could also be observed at the level of individual units, where the firing rates of single units would increase before, during, and after (depending on the unit) the burst onset time in both patients (**Fig. 4h**). Interestingly, between 25-55% of the individual units within different WaveMAP unit classifications demonstrated significant spike rate modulation relative to the burst onset while the other units did not vary significantly with the burst suppression. In addition, some units increased firing preceding the burst while others increased firing later in the burst, indicating separable response profiles (**Fig. 4h**). Together, these recordings appeared to provide richly detailed information about the spike-field relationship of neurons and their dynamics during physiologically relevant states.

## DISCUSSION

Here, we demonstrate that Neuropixels probes can be adapted for use in acute recordings of neuronal activity in the human cortex in an operating room environment. We found that, within minutes after inserting the Neuropixels probe into cortical tissue, we could observe up to 200 clear and stable units. Taking advantage of the spatiotemporal scales afforded by these probes, we could separate well-identifiable units based on waveform shapes (using duration and polarity), and we could also classify the waveforms using unbiased techniques via PCA with k-means clustering and, separately, using a novel nonlinear approach, WaveMAP ^55^. Similar waveform classes were found across participants using each of the three separate classification approaches, indicating there were consistencies between individuals. This waveform diversity presumably relates to both differences in cell types and differences in where electrical activity is recorded (e.g., near soma versus axons) ^55^.

Historically, single-channel extracellular recordings have led to the primary categorization of neurons into inhibitory vs. excitatory / pyramidal neuron classes. Since Cajal’s work, we have known that there are subclasses of these types based on location in the cortex, channel expression, morphology and more. It is possible that with this ultra high-density recording approach enabled by Neuropixels we can make strides towards associating electrophysiologically identified cell types with some of these detailed neuronal subtypes ^82–84^. Future work in both animal models and humans will be necessary to correlate functional differences in waveform morphology with essential cellular characteristics. In addition, even with this small sampling, we could find evidence of pairwise covariance between units. While such correlational analysis alone does not permit definitive elaboration of local circuit interconnections, it suggests that the high spatial sampling of the Neuropixels probe, especially if coupled to other sources of information such as detailed dual-patch slice work ^85^ and super-high resolution anatomical work ^86^ may provide detailed microcircuit maps of human cortex.

We also observed significant relationships between LFP bursts and unit spiking activity, with some units’ firing increasing even before the burst and others firing during and after. This possibly indicates a circuit-scale phenomenon that warrants further investigation as it may relate to the mechanism of burst-suppression in anesthesia ^72^. Further, we found some units would fire at and after an IID, indicating that the Neuropixels probe could capture clinically relevant activity in the LFP while sampling high resolution information about single cells, even acutely in the operating room and even in Pt.03’s recording where we “only” isolated a relatively fewer 29 units from this single probe.

Contrary to expectations based on previous reports from the primate literature, we did not observe a reliable bi-modal distribution of spike waveform duration. Historically, a bi-modal distribution has been used to identify putative pyramidal and inhibitory neurons (e.g. ^59^), but we did not find strong differences in firing rate for the RS versus FS units, as has been previously reported in the literature ^47, 50, 59, 60, 87^. While such a separation has been reported with Utah arrays in humans ^50, 87^, we saw no evidence of this distribution in the recordings reported here. One possible explanation is that the higher density of Neuropixels allows more sensitive neuron detection, which leads to recording a wider range of neurons with less bias towards large waveform action potentials. A second (not mutually exclusive) possibility is that this difference is due to the Neuropixels probe sampling across cortical layers, versus the more within-layer sampling of Utah arrays.

An important component of Neuropixels recordings in the operating room, and a consideration when examining activity across cortical layers, is that the range of possible probe angles and placement relative to gyri and sulci are both significantly increased by the flexibility of the small Neuropixels form factor in penetrating neural tissue from different angles (although approach trajectories are limited by the clinical considerations and access boundaries). In most cases, we could only approach the tissue with a limited set of angles (e.g. limited by the Burr hole) or penetrate a curved surface (with gyri more complex in humans compared to NHPs and mice), such that we may not be as perpendicular to the cortical layers as in NHP or mouse studies ^1, 3^.

An important advantage of the Neuropixel probes’ single-shank form factor is that it allows for considerable flexibility in the angle and direction of the insertion compared to a planar array like the Utah array ^17^. This additional flexibility of insertion configurations, however, introduces additional challenges for recording the spatial location and angular orientation of the probe. This requires additional measurement and documentation of these degrees of freedom, including: 1) mapping to whatever stereotactic placement system is used in 3D, 2) 3D reconstruction of the brain and Neuropixels probe relative to trajectories and photographs (from multiple angles, made possible with the use of Blender), and 3) the use of LFP (**Fig. 1**) and/or the use of different Neuropixels recording maps to define recording depth and boundaries. This approach could be additionally refined in future experiments by using the BrainLab software package (Munich, Germany) or other intraoperative visualization tools ^88^ to improve the mapping precision and fidelity.

For research with intraoperative time constraints, there is a high priority on being able to achieve high fidelity, stable recordings quickly. Previous experiences suggest that this may be hampered by cortical ‘stunning’ which results in few units being recorded for minutes or even longer after electrode insertion ^40^. With the Neuropixels probe, however, we were able to record large numbers of units within minutes of insertion. In two cases, we recorded more than 200 single units in a period of 5-10 minutes, while, in the third case, we recorded 29 units in a similar time span (10 minutes). The possible difference in the third participant was that the electrode was placed in the lateral temporal lobe to be resected for epilepsy, and the patient was deeply anesthetized (as indicated by the high level of suppression in the detected burst-suppression waves).

In addition to being able to record from large numbers of units, the Neuropixels arrangement allows for resolution of units and LFP at different cortical depths. It was clear that distinct types of units are distributed differently across the recording probe, with a possibility that we could also record backpropagating action potentials. Indeed, the dense sampling allows identifying many pairs of interacting neurons (i.e., those exhibiting a consistent temporal lead/lag). Previous reports of groups using Neuropixels to observe backpropagating action potentials, combined with the present results confirming the ability to super-sample action potentials across multiple electrodes, indicates that similar *in vivo* investigations of human action potential variants are possible ^2^. In addition, further analyses of the LFP alone could be mapped to sinks and sources using current source density techniques ^89–91^.

While our Neuropixels recordings of single and multi-unit activity in humans are promising, it is crucial to recognize that there are inherent limitations of this recording system for human neuroscience. In current configuration there is only a single, essentially linear array of electrodes allowing sampling along a single cortical column. Unlike in rodent and NHP experiments, widespread sampling is not yet available. Further, in some situations, data is sometimes obtained under anesthesia which will, of course, have an impact on the observed physiology, particularly as the spiking activity of individual neurons can stretch and shift through time due to anesthesia as shown in previous rodent Neuropixels studies ^71^. However, as our analyses demonstrated, this factor remains an exciting feature of this work since understanding the single unit activity mechanisms underlying anesthesia, as demonstrated in the burst-locked single unit activity during the anesthesia-induced burst suppression in two participants (**Fig. 4**), has broad prospective clinical implication. Future work with additional data can involve more detailed analyses of these relationships ^73^.

In addition, as expected, we encountered substantial challenges with the electrically noisy environment of the operating room. This required flexibility in the arrangement of the ground and reference electrodes for the recordings. Another major challenge is impact of brain movement on the recordings. Here, we were able to perform a post-hoc manual analysis to mitigate the artifact introduced by brain pulsation and have shared detailed instructions with code (see **Online Methods**). While this was sufficient to allow for accurate spike sorting for the units reported in this analysis, it is likely that many theoretically isolatable units with smaller signals were rejected or ignored due to the challenge of isolating small waveforms across drift events. We found that we could not rely on alignment approaches that only used spike rates^4^, since the movement artifact was considerably faster than the 1-2 seconds needed for published stabilization approaches^4, 48^. This is because the problem is not one of drift (i.e. slow displacement of the electrode) but of pulsation artifact due to the ejection of blood into the great vessels with each heartbeat. Our manual approach provided a reasonable fix for this problem but can likely be improved. To this end, we have provided open-source de-identified data which we hope will allow groups to develop/update automated alignment algorithms to work well with these new type of human electrophysiology data. Future work to automatically adjust for movement artifacts would ideally use the voltage fluctuations in the LFP, or a combination of LFP and spike rates, to align the channels and adjust for the movement.

Other methods currently under development to address this challenge aim to directly minimize the tissue motion during the recordings. Most of these approaches focus on mechanically stabilizing the cortex around the probe. Another approach would be to allow the recording electrodes to move with the brain, e.g. as developed by Neuralink ^92^ or with a flexible link isolating the probe from any positioning equipment or cables. We believe that further development of motion stabilization techniques -- in a manner which optimizes maximizing patient safety – is a top priority and will result in substantially more recorded neurons, each sampled and isolated with more consistency. Lastly, alternative electrode configurations, such as the linear column configuration used in the Neuropixels 2.0 probe ^4^ specifically to facilitate automated post-hoc motion correction, will likely help to address this challenge.

A further challenge to applying this technology in intraoperative studies is the necessary time constraints on the recordings. For patient safety, it is not appropriate to extend their time in the operating room significantly. Our current study limited extra time conservatively to ∼15 minutes, but this can be extended in the future, depending on the exact surgical context. For example, extensive language mapping in an awake tumor resection case may go on significantly longer. However, the short recordings could be a factor in the waveform shapes we observed (e.g., cell types that fire infrequently or only in certain contexts would be less likely to be observed). Studies in other species could indicate optimal recording stability is achieved at considerably longer recordings (45 minutes)^3^. While we cannot change this feature of our studies, we note it is a potential factor that affects our recordings. This open question presents an opportunity for comparable studies in NHPs or large animals to examine single units in those first moments of insertions, and then afterwards, in order to help us better contextualize human Neuropixels recordings.

Nevertheless, the fact that we obtained single unit-resolution recordings almost immediately upon insertion of the Neuropixels is highly encouraging, since it ensures that useful information can still be obtained in a short period of time. Indeed, the power of these acute tests in the OR using Neuropixels probes will be realized with experiments in which awake subjects engage with a task during the recording period. All such uses cases, however, necessarily build on a demonstration of the feasibility of acute recordings in clinical settings given the unique challenges in this environment. Perhaps most exciting is the potential for Neuropixels (or future variants) to accelerate clinical electrophysiology via dramatically increased channel counts relative to single- or low channel count probes. For example, simultaneously recording at multiple depths along a probe during deep brain stimulation targeting, instead of “searching” by moving the electrode through the brain ^9^ with single electrodes would reduce procedure time and potentially improve patient outcomes. Similarly, these probes allow for a detailed exploration of neuronal activity underlying epileptiform activity, the effects of anesthetic agents and the pathophysiology of brain tumors, all of which could benefit from the use of high resolution, microscale information on single neuron activity underlying these clinically relevant conditions ^30, 33, 45, 46^. Of course, there is a concern that sampling neural activity from a single cortical column will not allow us to identify the underlying neural circuits of cognition or pathophysiology. However, considerable work using laminar electrodes ^93, 94^ (**Fig. 3a**) and Utah arrays ^42, 44^ as well as single lowered electrodes as done during DBS surgeries ^32, 77^ have demonstrated that we can parse, and decode, complex cognitive processes and motor control even from a small area of cortex. In addition, recent work using higher resolution electrodes has revealed details previously inaccessible with current clinical technologies including microscale dynamics of epileptiform activity and travelling dynamics of interictal discharges ^24, 95^ and propagating wave dynamics across the hippocampus ^26^ with the promise that such thin film electrodes combined with Neuropixels arrays could open up an entire field of discovery, linking inter-cortical columnar dynamics as well as inter-laminar interactions. With these technical adaptations to the clinical domain as reported here and with the future possible addition of LFP-driven motion correction, gathering further data will necessitate asking essential, and exciting, questions surrounding neuronal dynamics including the independence of single neurons relative to populations ^68, 69^ or relative to broad LFP activity such as travelling waves ^70^. Indeed, substantial additional research avenues will be opened by technological advances which enable a chronic form of the system.

In summary, the Neuropixels approach suggests a pathway forward for increasingly sophisticated and detailed explorations of the cellular-scale code underlying higher order cognitive function in multiple areas of the human brain, as well generating a deeper understanding of the dynamics of clinically relevant neural activity. It is also a step towards developing high channel count chronic neural interfaces for human use which may accelerate and expand the therapeutic possibilities of brain-computer interfaces ^96^.

## Extended Data / Supplementary Information

### Online Methods

**Supplemental Table 1. Distribution and numbers of cases and results as well as reasons for data exclusion and information per participant and recording**

**Supplemental Figure 1. Recording challenges and lessons learned.**

**Supplemental Figure 2. Putative Neuropixels probe location based on photographic evidence and identification of intracranial access to the cortical surface**

**Supplemental Figure 3: Realigning the data relative to heartbeat-induced movement artifact.**

**Supplemental Figure 4. Cross correlation between the interpolated sorted units and the raw data**

**Supplemental Figure 5. Example complex waveforms for six different units (each color-coded set of waveforms) across the data set.**

**Supplemental Figure 6. Waveform measures.**

**Supplemental Figure 7. Spatial waveforms measures.**

**Supplemental Figure 8. PCA with k-means clustering**

**Supplemental Figure 9. Waveform Features of Units Clustered with WaveMAP.**

**Supplemental Figure 10. Single units through time and correlation relationships between units.**

**Supplemental Video 1. Raw and Interpolated Neuropixels data.**

## Acknowledgements

We would like to thank Yangling Chou, Aaron Tripp, Fausto Minidio, Alex Zhang, and Alexandra O’Donnell for help in data collection. We would like to especially thank the patients for their willingness to participate in this research. This research was supported by the ECOR and K24-NS088568 (to SSC) and the Tiny Blue Dot Foundation (to SSC and ACP) and NIH grant U01NS121616 (to ZMW). This research was also supported by NIH NINDS BRAIN R01NS11662301 (to KVS), NIH NIDCD R01DC01403406 (to KVS), the Simons Foundation 543045 (to KVS) and the Howard Hughes Medical Institute at Stanford University (to KVS, SDS, EMT). SDS was supported by the A. P. Giannini Foundation, Wu Tsai Neurosciences Institute Interdisciplinary Scholars Fellowship and holds a Career Award at the Scientific Interface from the Burroughs Wellcome Fund. EMT is additionally funded by the Brain and Behavior Research Foundation and the Grossman Institute. The views and conclusions contained in this document are those of the authors and do not represent the official policies, either expressed or implied, of the funding sources.

## Author Contributions

The experiment was conceived by SSC, YK, ACP, ZW, KVS, LRH, EMT, and SDS. ZW and RMR performed the surgeries and placed the arrays while ACP, MM, AK, and YK collected the data and did first pass analysis. BD, MW, and EMT conceived of and advanced the production of the thicker custom Neuropixels probes used in the study. All the authors edited the manuscript.

## METHODS (COMPLETE ONLINE VERSION)

### Patients & clinical/research electrode placement

All patients voluntarily participated after informed consent according to guidelines as monitored by the Partners Institutional Review Board (IRB) Massachusetts General Hospital (MGH). Participants were informed that participation in the experiment would not alter their clinical treatment in any way, and that they could withdraw at any time without jeopardizing their clinical care. Recordings in the operating room were acquired with 9 participants (mean= 59 years old, ranging from 34 to 75; 7 female; **Supplemental Table 1**) who were already scheduled for a craniotomy for concurrent clinical intraoperative neurophysiological monitoring or testing for mapping motor, language, and sensory regions and removal of tissue as a result of tumor or epilepsy or undergo intra-operative neurophysiology as part of their planned deep brain stimulator (DBS) placement ^1–4^. Prior to inserting the Neuropixels probe, a small superficial incision in the pia was done using an arachnoid surgical knife. The Neuropixels probe was inserted through this incision. Recordings were referenced to sterile ground and recording reference needle electrodes (Medtronic) placed in nearby muscle tissue (often scalp) as deemed safe by the neurosurgical team though a series of tests ground and reference tests were performed to identify the ideal combinations of ground and reference options, listed below (**Supplemental Table 1**).

Following the surgery, the preoperative T1-weighted MRI was used to generate a 3D surface brain map using FreeSurfer scripts ^5–7^ (http://surfer.nmr.mgh.harvard.edu). Images obtained during surgery, locations as indicated using Brainlab (Brainlab, Inc.), and photographs captured during the surgery were aligned to the 3D reconstructions using Blender software (https://www.blender.org/) and MMVT ^7–9^ (**Supplemental Figure 2**). For the two DBS cases, there are physical limits imposed by the three-dimensional shape and location of the burr hole as well as the size of the Neuropixels probe and the headstage. As such, we took the postoperative CT after the DBS leads were implanted (after the surgery) and overlaid it with the preoperative MRI using Mango ^10–12^. We then used these coordinates and sizes to reconstruct the to-scale burr hole and DBS lead trajectories and mapped them to the participants’ brains reconstructed using MMVT and Freesurfer ^5–9^, and then placed the Neuropixels probe in Blender to the best approximation based on all this information. For the open craniotomy case, the reconstruction involved projecting the surgical image onto the patient’s reconstructed brain using Blender and then placing a 3D model of the Neuropixels probe on that location similar to other coregistration approaches ^1, 7, 9, 13^ (**Supplemental Figure 2**). Angles were calculated from photographs taken during the surgery as well as trajectories limited by the location and angle of the burr hole for DBS surgery.

Separately, to compare the relative sizes of the Neuropixels probe contacts relative to the reconstructed pyramidal neurons and inhibitory interneurons from NeuroMorpho.Org ^14–18^. The process involved importing the .swc files from NeuroMorpho.Org into MATLAB (Mathworks, Natick, MA), converting the data into a surface and then exporting the .stl files which we then imported into Blender. After manual adjustments, the reconstructed neurons were then placed next to the to-scale 3D reconstructed electrodes in Blender (**Fig. 3a, c**).

### Neuropixels recordings, data collection & analysis

Neuropixels probes (NP v 1.0, version S, IMEC) sterilized with Ethylene Oxide (BioSeal) were connected to a 3B2 IMEC headstage wrapped in a sterile plastic bag and sealed using TegaDerm (3M) to keep the field sterile. Neuropixels probes (NP v 1.0-S, IMEC) include an electrode shank (width: 70µm, length: 10 mm, thickness: 100µm) of 960 total sites laid out in a checkerboard pattern with contacts at ∼18 µm site to site distances (16 µm (column), 20 µm (row); ^11^). Handling of the electrodes and the headstage from outside the sterile bag was all performed in sterile conditions in the operating room. The headstage was connected via a multiplexed cable to a PXIe acquisition module card (IMEC), installed into a PXIe Chassis (PXIe-1071 chassis, National Instruments). For the Neuropixels-1.0 probes as used in this study, the linear dynamic range of the Neuropixels amplifier is 10 mVpp. This range is digitized using a 10 bits Analog to Digital conversion ^19^. All Neuropixels recordings were performed using SpikeGLX (http://billkarsh.github.io/SpikeGLX/) on a computer connected to the PXIe acquisition module recording the action potential band (AP, band-pass filtered from 0.3-10 kHz) sampled at 30 kHz and a local field potential band (LFP, band-pass filtered from 0.5-500 Hz), sampled at 2.5 kHz ^20–22^. Since these Neuropixels probes enable 384 recording channels which can then be used to address 960 electrodes across the probe shank, we tested different electrode maps which allowed us to record different portions of the probe. One map allowed for recording the lower portion of the probe (the most distal channels). A second map allowed for recording two rows along the entire length of the electrode. This map was used to identify the depth of the electrode in the cortex and we switched to the distal tip map (short map) for the main recording. A final map allowed for recording in a series of tetrode locations, skipping rows to distribute recordings along the entire length of the probe.

As the Neuropixels probe is stably rigid with regard to the channels relative to one another, we could estimate depth of the sites in the tissue along the electrode using the LFP signal. When some channels are outside the tissue, we found the LFP signal was relatively noisy and did not show differences between channels when outside of tissue (which we validated in separate tests in mice and in saline, data not shown). This electrophysiological marker, however, provided an upper boundary limit of how deep the electrode was in the tissue relative to the measurements on the electrode itself resulting in a depth relative to this upper boundary. However, we did not identify the depth as cortical depth as the Neuropixels probe could be inserted at different angles relative to the sulci and gyri (**Supplemental Fig. 2**).

Synchronization was performed through two different approaches. TTL triggers via a parallel port produced either during a task via MATLAB or custom code from a separate computer were sent to both the National Instruments and IMEC recording systems, via a parallel port system. In addition, we used the TTL output to send the synchronization trigger via the SMA input to the IMEC PXIe acquisition module card to allow for added synchronizing triggers which were also recorded on an additional breakout analog and digital input/output board (BNC-2110, National Instruments) connected via a PXIe board (PXIe-6341 module, National Instruments). The TTL triggers were produced either during a task via MATLAB or custom code on the task computer.

### Recording challenges and lessons learned

Five main challenges were faced when performing these recordings: 1) sterilization and maintaining a sterile field and conditions; 2) electrode fracture and disconnects; 3) decreasing noise in the recordings through referencing; 4) external sources of noise; 5) mechanical stabilization (**Fig. 1**; **Supplemental Fig. 1**).

#### Sterilization and maintaining a sterile field

To ensure we could use the Neuropixels probes in the OR, we worked with BioSeal (Placentia, CA) and sent them a sample of 25 Neuropixels probes. BioSeal took the samples through a validation process to determine that ethylene oxide (EtO) could be used to sterilize the Neuropixels probes. We also tested whether working Neuropixels probes were operational before and after sterilization. An important part of the process was identifying safe sterile packaging for sterilization and transport. We found we could place the probes sideways inside a slightly modified EtO-safe sterile container (SteriBest Trays, Sterilization Instrument Tray, Instrument Tray Sizes (inches):Base, Lid, Mat 6×2.5x.75], item#A-CP614, from Duraline Biosystem; **Fig. 1a**; **Supplemental Fig. 1g**; **Supplemental Table 1**). When received the boxes, we clipped protruding silicone nubs in an area of 3 cm x 3 cm on one side of the box as well as a few silicone nubs on the other end of the box. We found that we could package and safely ship and handle the Neuropixel probe cross-country by weaving the Neuropixels ribbon cable around the vertical silicone nubs in the sterilization containers with the Neuropixels probe and headstage perpendicular to the base of the box. We performed several tests to demonstrate the probe consistently survived this shipment approach, including before and after sterilization. Before shipping for sterilization, we soldered on a 10 cm long male touchproof cable (the white cable in **Fig. 1a**, **b** and **d**) to the reference side of the Neuropixels probe. In addition, we labelled the lid of the box to track individual probes. The validation of 25 probes performed by BioSeal was done with the Neuropixels probes in this configuration and with this specific SteriBest Tray packaging (including the added touchproof connection cable). Once shipped to Bioseal packed in bubble wrap, the company would return the probes in their sterilization boxes sealed in approved packaging. We have found this approach kept the electrodes intact and tracked throughout transport and sterilization.

#### Electrode fractures and disconnects

We had instances of electrode fracture (N=2), both of which were with the thinner Neuropixels 1.0 probes (thickness: 25µm, width: 70µm, length: 10 mm). We then switched to a thicker custom Neuropixels 1.0-S probe (thickness: 100µm width: 70 µm, length: 10 mm) for the remaining recordings and, of the 7 uses of thick probes, we only had one instance of electrode fracture. In each instance, we documented whether the probes were intact afterward both via the SpikeGLX software and through thorough photograph documentation. In the three probes which were fractured, we were able to photograph the pieces to reconstruct the entire probe to validate probe recovery. In the remaining probes, the photographs after the case confirmed the electrodes were fully intact after the case. In addition, in the intact probes after the case, software check via SpikeGLX involved a hardware check indicating the probes were intact and fully functioning.

We ensured insertion of the electrodes were done under direct visual guidance for the entire process with loupe magnification therefore allowing us to identify a mechanical break if it occurred. Throughout the entire process, the real-time recordings and impedance testing during placement and insertion provided real-time confirmation that the electrode had any stress or strain (which resulted in an error message in the software). If at any point, a fracture/disconnection was noted, the electrode advancement was halted, and the fractured electrode removed. Importantly, though, and as communicated to the participants, only regions that were planned for resection or cortisectomy were implanted. These areas were therefore already planned for removal meaning that, even in the event of a retained electrode, the tissue together with electrode can be safely/ethically removed. Finally, Neuropixels probes are radiopaque meaning that they can be identified by imaging if needed. Importantly, all these considerations were always discussed with the participants prior to study consent. There were no instances of retained electrode components in our studies

We had investigated the use of a temporary stabilization using TISSEEL (Baxter) as has been used with single fine wire recordings ^3^, but we found that, with the use of TISSEEL, we could not visualize where the Neuropixels probe was inserted which was problematic as we were aiming for the small incision on the pial surface. In addition, we found the sealant placed additional stress on the array shank, therefore increasing the risk of fracture.

#### Decreasing noise using referencing and grounding

Even though we had tested the Neuropixels probe as well as had considerable experience in using Neuropixels in NHPs which informed how we built our electrophysiological system ^22^, we found moving the Neuropixels recordings into the human OR was made much more difficult with considerable added noise compared to any of the other testing settings. In the first four tests, we followed the original recommendations to tie the reference to the ground on the Neuropixels probe which degraded the signal considerably in the OR (**Supplemental Fig. 1a**). The signal was substantially improved by separating the ground and reference on the Neuropixels probe, with a single Medtronic sterile wire connected to the reference placed in the scalp and a separate wire attached to the ground and also placed in the scalp as deemed safe by the neurosurgical team. Improving the signal also involved tying the patient ground to the recording ground the patient to the recording via a BOVIE pad (Clearwater, FL) connected to the grounding BNC on the NIDAQ board used for the Neuropixels system. Placing the grounding lead into saline or CSF degraded the signal by saturating the LFP and increasing noise in the system.

#### External sources of noise

Changing the reference from the external reference in the software (using SpikeGLX) to the internal reference also increased noise significantly (**Supplemental Fig. 1c**). We also discovered an external source of noise was the wall-powered anesthesia IV pump (as is commonly used during patient transport) which, when unplugged and operating on battery, would decrease the physiological noise. Finally, we did a series of tests to determine if other signals added sources of noise and we did not find an effect of the BOVIE cautery machine, the ROSA robot, the lights or other machines in the room.

#### Mechanical stabilization

Two separate stabilization approaches were tested. One approach involved the patients receiving DBS implantations at MGH, who normally also undergo standardized micro-electrode recording to optimize anatomical targeting ^4, 23^, Neuropixels probes were inserted in the same locations as the microelectrodes that traverse the dorsal lateral surface of the prefrontal cortex on the way to the target nucleus, offering a brief chance to study neuronal dynamics in the dlPFC and not perturbing the planned operative approach nor alter clinical care ^2–4, 23–25^. Three cannulae were placed in a manipulator (AlphaOmega Engineering, Nazareth, Israel) and the Neuropixels probe was attached to the cannulae using SteriStrips (3M™ Steri-Strip™ Reinforced Adhesive Skin Closures). The manipulator was attached to the ROSA ONE® Brain (Zimmer Biomet) arm. The Neuropixels probe was put over the burr hole by the ROSA robot arm. ROSA was then used to move the probe insert the probe using fine millimeter steps, with some adjustment possible using the AlphaOmega micromanipulator. The second approach involved securing the Neuropixels probe to a sterile syringe which was then held by a 3-axis micromanipulator built for Utah array placement (BlackRock, Salt Lake City, UT) which was attached to a Greenberg retractor. The Neuropixels probe was in place and lowered using the micromanipulator.

### Compensation for tissue movement and electrode alignment through time

We found clear evidence of vertical tissue movement relative to the Neuropixels probe in the local field potential (LFP) recordings (**Supplemental Fig. 2**). To confirm that this was due to movement of the tissue as well as effects of heartbeat, we aligned the movement artifact to the heartbeat in time (this was possible thanks to audio tracking of the EKG in 2 participants’ cases). We found the movement roughly matched this tracking. To confirm that the manual tracking could match the movement of the brain relative to the electrode, we performed tissue-level tracking of the video recordings of the case and found we could align the filmed movement of the brain pumping relative to the electrode, which was well visualized in the LFP band across channels as tracked through time (**Supplemental Fig. 2b**). We tested several approaches to address this movement and correct for the alignment, including the Kilosort 3.0 drift adjustments and estimation (https://github.com/MouseLand/Kilosort) and spike time-informed alignment approaches (https://github.com/evarol/NeuropixelsRegistration). We chose to use the LFP-informed manual tracking as it was better-resolved in the time domain since the dynamic range of LFP allowed for per time step (0.0004 sec) alignment and interpolation. In contrast, the automatic approach depended on firing rate and arrival of spikes, which were sparse (**Fig. 1e-f**).

#### Manual tracking of movement using LFP signals

The signal was first extracted from the binary files into local field potential (LFP, <500 Hz filtered data, sampled at 2500 Hz) and action potential (AP, >500 Hz filtered data, sampled at 30000 Hz) from SpikeGLX using MATLAB and available preprocessing code (https://billkarsh.github.io/SpikeGLX/). We inspected the data visually as well as examined the timeline of the recording to reject noisy time ranges (such as during insertion.) We then further examined the voltage deflections in the LFP for a prominent, bounded deflection in the voltage where we observed the voltage values shifting in unison (**Supplemental Figure 2**) which was consistently present throughout the recording (blue or red bands in **Supplemental Figure 2**). We attempted to use a number of algorithms to detect these shifts, but the multiple changes present (heartrate, slow and mid-range drifting, and other shifts) were not effectively tracked by these algorithms. Instead, to capture the displacement in the movement bands, we imported the LFP voltage as an .stl file from MATLAB into Blender (https://www.blender.org/), a three-dimensional animation program which allowed for easier manual tracing compared to MATLAB. Using the surface voltage and the Grease Pencil feature, we traced the shifting band of negatively deflecting LFP throughout the recording at a resolution of 500 Hz. The line produced then was exported as a .csv file and imported into MATLAB, where it was compared with the LFP at higher resolution to check whether the manual tracing matched the LFP displacement (**Supplemental Figure 2a**). This traced line information was upsampled to 2500 Hz to match the sampling frequency of the LFP channels (interp1, ‘makima’).

#### Preprocessing AP recordings

Once we had the LFP baseline to track probe movement through time, we then applied analyses to the AP sampled band. To account for differences in the channels before aligning the data (as channels can have differences in impedance), we first detrended data (which removes best fitted line to each channel), calculated the median, and subtracted it from all channels. We then normalized the voltage signal across channels by multiplying each channel’s voltage time series by a normalization factor where *Normalized data = Channel signal * (1/std) * 600*. In this case, the *std* was the standard deviation of channel data without outliers, particularly epochs which were relatively quiet. We defined outliers as elements which were more than 1.5 interquartile ranges above the upper quartile or below the lower quartile of the distribution of voltage signals. Finally, we chose the value of 600 in the normalization to allow us to scale the data up to an int16 format for improved data resolution.

#### Alignment and interpolation of AP channels for manual registration

To then re-align the AP channel data so as to offset the movement artifact, we upsampled the traced line to 30KHz to match the AP sampling rate (interp1, ‘makima’). We then, for each time bin, applied a spatial interpolation between channels vertically in two columns of the Neuropixels recording, resulting in a vertical spatial resolution of 1um. (**Supplemental Figure 2**). These steps resulted in a large, high resolution interpolated matrix that we could then follow through time. This let us compensate for the movement effects by resampling the voltage in space (**Supplemental Figure 2**) based on the manually registered movement trajectory described in “Manual Tracking of Movement using LFP signals”.

Specifically, for each time bin, we shifted the vertical channels vector up or down according the upsampled traced line, resulting in >450 ‘virtual channels’ that each contained voltage information putatively from a specific brain location. Finally, since the virtual channels on both ends (top and bottom of the shank) contained only partial data (due to brain movement relative to the electrode), we selected a subset of 384 virtual channels that contained the most continuous information throughout the recording (and did not shift channels into the edge), which could be inferred from the average channel offset.

### Unit isolation and clustering

Single unit sorting was performed using Kilosort 3.0 ^26^ (https://github.com/MouseLand/Kilosort) as well as Phy (https://github.com/cortex-lab/phy) and then manually curated using in-house MATLAB code to visually inspect the template as well as the waveforms assigned to each cluster. The Kilosort 3.0 parameters included: Nblocks = 0 – as no additional registration was needed according to spiking activity after the manual registration; Threshold [10, 11] to be more strict in our detections (initial values were [9,9] which resulted in ∼800 units for Pt. 02). Clusters were merged in Phy if the templates were similar between clusters, the spatial spread of waveforms were highly similar and overlapping, and cross-correlations of the event times indicated high levels of correlation. To further process and shift each individual waveform to correct for Kilosort3 misalignment, we also calculated the cross correlation of individual waveforms with the cluster template and adjusted waveforms according to location of maximal voltage value per waveform in the sampled time.

### Waveform feature analyses and classification

Clusters were then separated into single units and multi-unit activity (MUA). Units were classified as MUA if there was a mixture of distinct waveforms (examined in Phy) as well as a complicated (and abnormal) autocorrelogram. For all units, we then measured the spike duration, halfwidth, peak-trough ratio, repolarization slope, recovery slope, and amplitude measures (**Fig. 2**; adapted from ^21^; https://github.com/jiaxx/waveform_classification). Further, we applied the spatial spread and velocity measures to each cluster to identify whether we could observe evidence for backpropagating action potentials or other unique spatial dynamics (**Fig. 2**; adapted from ^21^).

We used three different classification approaches to group the units. First, using a standard approach, units were grouped into regular spiking (RS), fast spiking (FS), positive spikes (PS) classifications based on the spike waveform duration (valley-to-peak) of the largest peak across channels per unit ^27–32^. The ranges for each classification were as follows: negative going peaks included FS (duration <0.3 ms) and RS (duration>0.3 ms) and positive spikes (PS). Second, we applied principal components analyses on the first six channels per unit and clustered these average waveforms using k-means clustering (squared Euclidean distance, 1000 replicates, 1000 maximum iterations) into 7 clusters based on the separability of the clusters (using silhouette) and how clean the resulting clusters were. Finally, we used a novel non-linear method, WaveMAP, which took into account the spatial and temporal waveform characteristics while separating out differences in the waveforms ^24^. WaveMAP includes a combination of dimensionality reduction with Universal Manifold Approximation and Projection (UMAP) combined with Louvain clustering to identify clusters in the data set ^33^. We then compared the waveform features across these different classifications.

### Local field potential analyses

Custom MATLAB code (version R2020a) in combination with open source code from the Fieldtrip toolbox (^34^; http://www.fieldtriptoolbox.org/).

### Burst suppression ratio measurement

The burst suppression ratio (BSR) was computed using an automated method ^35, 36^ (https://github.com/drasros/bs_detector_icueeg). After averaging the LFP across all channels, this method then labels each time sample as either burst or suppression. Briefly, the method uses the previous data with each channel and applies the following equations:

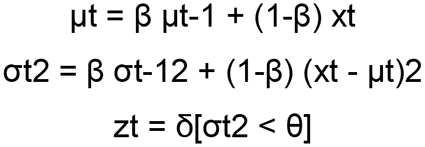

Where xt is the value of the normalized signal of one channel at time t, µt and σt2 are current values of the recursively estimated local mean and variance, respectively. Finally, zt is an indicator function that labels each data point as either a burst (0) or suppression (1). The value of β determines the balance between the effect of recent and past data set based on previously trained data ^35^. The classification threshold θ (i.e., the value above which a data point should be classified as burst) was adjusted to evaluate our dataset visually with values of θ = 50, 100, 150, and 200 and was informed by the input from two experts who reviewed selected intervals to identify burst and suppression using each possible theta value. The value of θ = 200 was selected to reliably identify burst and suppression induced by general anesthesia. The burst suppression ratio for each recording in an anesthetized patient (N=2) was evaluated as the proportion of suppression-labeled samples in a moving window (1 s duration, no overlap).

### Inter-ictal epileptiform discharges

In one case, the Neuropixels electrode was inserted into the lateral temporal lobe before tissue resection for epilepsy. As we could identify interictal epileptiform discharges (IIDs) in the LFP, we applied both an automatic and a visual detection approach to verify the timing and location of the IIDs in the Neuropixels recording. For automatic detection, we averaged the LFP across channels and applied the algorithm of ^37^, version v21, default settings except -h at 60 and -k1 at 7 to increase the threshold for detection; http://isarg.fel.cvut.cz), which adaptively models distributions of signal envelopes to discriminate IIDs from LFP ^37^. In addition, a trained and experienced epileptologist (SSC) examined the average LFPs and confirmed the timing of the detected IIDs. This two-step process was necessary as the burst suppression from the anesthesia produced waveforms which could obscure the IIDs. For several analyses, the single unit spike times were then aligned relative to the peaks of the IIDs.

### Statistical analysis

All statistical comparisons were performed using non-parametric measures, so we did not test for normality. Multiple comparisons tests were performed using the Kruskal–Wallis test for non-equivalence of multiple medians followed post hoc Tukey-Kramer method to identify statistically separable groups. For comparisons between individual medians, we used the Wilcoxon rank-sum test (two-sided). We corrected by adjusting the target p-value (0.05) with a Bonferroni correction for the number of comparisons being done.

### Data availability

The data will be available for download at Dryad (https://datadryad.org/stash) upon publication.

### Code availability

Open source acquisition software, SpikeGLX (http://billkarsh.github.io/SpikeGLX/) and record the neural data. Single unit sorting was performed using Kilosort 3.0 ^26^(https://github.com/MouseLand/Kilosort) as well as Phy2 (https://github.com/cortex-lab/phy) Custom Matlab code (version R2020a) and python code in combination with open source code from the Fieldtrip toolbox (http://www.fieldtriptoolbox.org/) was used for the majority of the analyses with some code involving manual alignment available on Github (https://github.com/Center-For-Neurotechnology/CorticalNeuropixelProcessingPipeline). The burst suppression ratio (BSR) was computed using an automated method ^35, 36^ (https://github.com/drasros/bs_detector_icueeg). Reconstruction of electrode locations and the manual tracing was done using the open source, free software Blender (https://www.blender.org/).

