## Supplemental Data for "Large-scale neural recordings with single-cell resolution in human cortex using high-density Neuropixels probes"

**Supplemental Figures and Extended Data for: Large-scale neural recordings with single-cell resolution in human cortex using high-density Neuropixels probes**


### AUTHORS

Angelique C. Paulk[1], Yoav Kfir[2], Arjun Khanna[2], Martina Mustroph[2], Eric M. Trautmann[3], Dan J. Soper[1], Sergey D. Stavisky[4], Marleen Welkenhuysen[5], Barundeb Dutta[5], Krishna V. Shenoy[6], Leigh R. Hochberg[1,7], R. Mark Richardson[2], Ziv M. Williams[2]*, and Sydney S. Cash[1]*

### AUTHOR INFORMATION

[1] Center for Neurotechnology and Neurorecovery, Department of Neurology, Massachusetts General Hospital, Boston, MA, USA; Department of Neurology, Harvard Medical School, Boston, MA, USA.

[2]Dept. of Neurosurgery, Harvard Medical School and Massachusetts General Hospital, Boston, MA, USA.

[3]Zuckerman Institute, Columbia University, New York, NY; Department of Electrical Engineering, Stanford University, Stanford, CA, USA; Wu Tsai Neurosciences Institute, Bio-X Program, Stanford University, Stanford, CA, USA; Howard Hughes Medical Institute at Stanford University, Stanford, CA, USA.

[4]Department of Neurosurgery, Stanford University, Stanford, CA, USA; Department of Electrical Engineering, Stanford University, Stanford, CA, USA; Wu Tsai Neurosciences Institute, Bio-X Program, Stanford University, Stanford, CA, USA.

[5]IMEC, Leuven, Belgium.

[6]Department of Electrical Engineering, Stanford University, Stanford, CA, USA; Department of Bioengineering, Stanford University, Stanford, CA, USA; Department of Neurobiology, Stanford University, Stanford, CA, USA; Howard Hughes Medical Institute at Stanford University, Stanford, CA, USA; Wu Tsai Neurosciences Institute, Bio-X Program, Stanford University, Stanford, CA, USA.

[7]VA RR&D Center for Neurorestoration and Neurotechnology, Rehabilitation R&D Service, Providence VA Medical Center, Providence, RI, USA; School of Engineering and Carney Institute for Brain Science, Brown University, Providence, RI, USA.

* Co-senior authors

### CORRESPONDENCE

Angelique C. Paulk, Sydney S. Cash and Ziv M. Williams

*Mail:* 55 Fruit St. Boston, MA 02114

**Number of Figures:** 4
**Number of Tables:** 0
**Number of Extended Data Figures and tables:** 11
**Number of Supplementary Videos:** 1

**Data availability**

The majority of the data that support the findings of this study are available from the corresponding author upon reasonable request, though a subset of data will be available for download at Dryad (https://datadryad.org/stash) upon publication.

**Code availability**

Open source acquisition software, SpikeGLX (http://billkarsh.github.io/SpikeGLX/) and record the neural data. Single unit sorting was performed using Kilosort 3.0 [26](https://github.com/MouseLand/Kilosort) as well as Phy2 (https://github.com/cortex-lab/phy ) Custom Matlab code (version R2020a) and python code in combination with open source code from the Fieldtrip toolbox (http://www.fieldtriptoolbox.org/ ) was used for the majority of the analyses with some code involving manual alignment available on Github (https://github.com/Center-For-Neurotechnology/CorticalNeuropixelProcessingPipeline). The burst suppression ratio (BSR) was computed using an automated method [35,36] (https://github.com/drasros/bs_detector_icueeg). Reconstruction of electrode locations and the manual tracing was done using the open source, free software Blender (https://www.blender.org/ ).

**METHODS REFERENCES**

**Extended Data Figures**

**Supplemental Table 1. Distribution and numbers of cases and results as well as reasons for data exclusion and information per participant and recording**

| Participant | Procedure | Location | Type | Recording and number of units | Reason for exclusion of data |
|---|---|---|---|---|---|
| | left anterior frontal lobe tumor removal, MAC and awake | Left anterior frontal lobe | Thin | No recording | Electrode fractured and recovered |
| | DBS implant, MAC and awake | Left dorsolateral prefrontal lobe | Thin | Short recording, but some fracture during recording | Electrode fractured and recovered |
| | DBS implant, MAC and awake | Left dorsolateral prefrontal lobe | Thick | Recording, but noise was significant | Electrode intact, noise considerable |
| | DBS implant, MAC and awake | Left dorsolateral prefrontal lobe | Thick | Recording, but noise was significant | Electrode intact, noise considerable |
| Pt. 01 | DBS implant, generalized anesthesia, not awake | Right dorsolateral prefrontal lobe | Thick | 262 total clusters, 202 single units, 60 MUA clusters | None, Electrode intact and recovered |
| Pt. 02 | DBS implant, MAC and awake | Left dorsolateral prefrontal lobe | Thick | 312 total clusters, 178 single units, 134 MUA clusters | None, Electrode intact and recovered |
| | left anterior temporal lobectomy, MAC and awake | Left anterior temporal lobe | Thick | Recording, but noise was significant | Electrode intact, but considerable noise |
| Pt. 03 | left anterior temporal lobectomy, generalized anesthesia, not awake | Left anterior temporal lobe | Thick | 29 total clusters, 19 single units, 10 MUA clusters | None, Electrode intact and recovered |
| | DBS implant, MAC and awake | Left dorsolateral prefrontal lobe | Thick | Recording, but noise was significant | Electrode intact, but considerable noise |

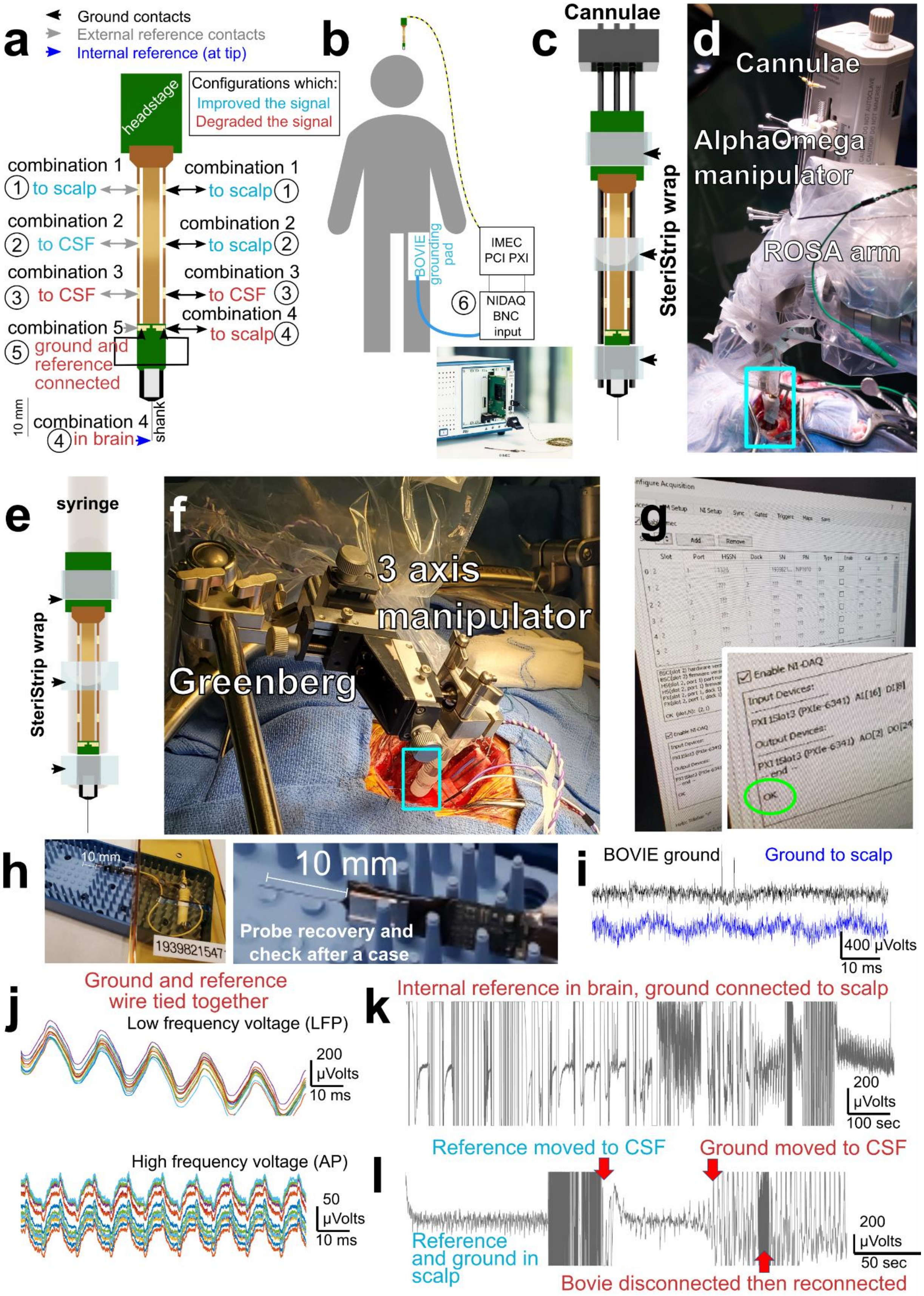
a
Ground contacts
External reference contacts
Internal reference (at tip)
Configurations which:
Improved the signal
Degraded the signal
headstage
combination 1
1 to scalp
combination 1
to scalp 1
combination 2
2 to CSF
combination 2
to scalp 2
combination 3
3 to CSF
combination 3
to CSF 3
combination 4
to scalp 4
combination 5
5 ground and reference connected
10 mm
combination 4
4 in brain
shank
b
BOVIE grounding pad
IMEC PCI PXI
6 NIDAQ BNC input
c
Cannulae
SteriStrip wrap
d
Cannulae
AlphaOmega manipulator
ROSA arm
e
syringe
SteriStrip wrap
f
3 axis manipulator
Greenberg
g
Enable NI-DAQ
OK
h
10 mm
1939821547
10 mm
Probe recovery and check after a case
i
BOVIE ground
Ground to scalp
400 µVolts
10 ms
j
Ground and reference wire tied together
Low frequency voltage (LFP)
200 µVolts
10 ms
High frequency voltage (AP)
50 µVolts
10 ms
k
Internal reference in brain, ground connected to scalp
200 µVolts
100 sec
l
Reference moved to CSF
Ground moved to CSF
Reference and ground in scalp
200 µVolts
50 sec
Bovie disconnected then reconnected

**Supplemental Figure 1: Recording challenges and lessons learned. a.** Diagram of the different combinations of ground and referencing and connections that improved the electrophysiological signal or degraded the signal with the indicated ground and reference contacts (including the external and internal reference). Each circled number indicates a combination tested. A major step in reducing noise levels was to separate the ground and reference, with a single separate wire going to the grounding pads on one side and another wire going to a grounding pad on the other side of the Neuropixels probe, pictured here. We used the external reference tied to a sterile MedTronic grounding wire with a needle which, when placed in the scalp or CSF, improved the signal and reduced 60 Hz noise. Common 60 Hz noise and other noise decreased significantly if the anesthesia IV pump was unplugged from the wall and was run on battery during the recordings. Otherwise, turning off lights or other sources of noise (BOVIE cautery machine, AlphaOmega recording system, etc.) had no noticeable effect on noise. **b**. Diagram of the connections relative to the participant including the BOVIE pad placed on the leg connected to the IMEC ground via a BNC cable to the NIDAQ PCI chassis BNC. **c**. Securing the wires and the probe to the cannulae attached to the AlphaOmega manipulator with Tegaderm and Steristrips improved the stability. **d**. Mechanical stabilization of the probe involved two options, one using the ROSA robot combined with an AlphaOmega manipulator with the Neuropixels probe secured to cannulae. **e**. The second option involved securing the Neuropixels probe to a sterile syringe using Steristrips. **f**. Then, the syringe is attached to a 3-axis manipulator mounted on a Greenberg retractor over the craniotomy. **g**. Every Neuropixels probe is documented and checked several times during and after the procedure both via software (in the OpenEphys or SpikeGLX software) and a quick saline test is done in the side sterile table. **h**. Each probe is photographed before, during, and afterward in the OR to determine if they are intact. Regular checks of the connections are done as the electrode is being moved into place in the sterile field. The electrode is also checked and photographed afterward for whether it is intact and connected. **i**. In two cases, 60 Hz noise was reduced by tying a ground to a spare BOVIE pad placed on the thigh of the patient under the sterile drape (as shown in **b**), with the traces shown with the lower left LFP trace showing the signal grounded to the BOVIE versus grounded to a scalp needle electrode. **j**. Examples of considerable noise in the LFP and high frequency range (AP range) when crossing the ground and reference together. **k**. We did test using the internal reference on the Neuropixels probe and found the noise increased significantly in the two cases we attempted the switch. **l**. The placement of the sterile ground and reference leads made a difference. Ground and reference in the scalp had an improved signal. Placing the ground (but not the reference) in the saline in the craniotomy caused the LFP signal to saturate and degraded the signal.

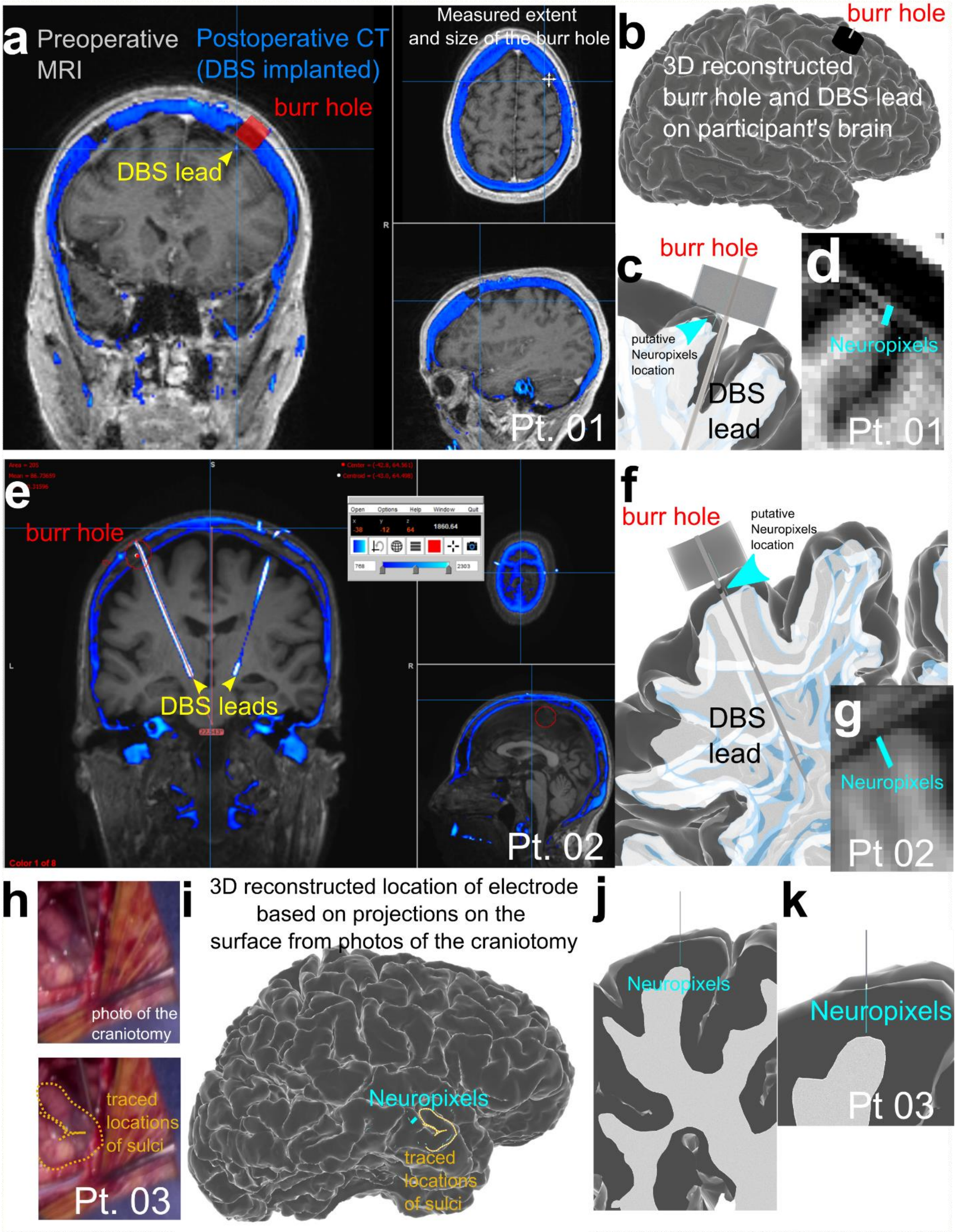


**Supplemental Figure 2**. **Putative Neuropixels probe location based on photographic evidence and identification of intracranial access to the cortical surface.** a. preoperative

MRI (Pt. 01) (displayed with Mango software[10–12]) with an overlay of the postoperative CT including the DBS leads and burr hole used to implant the DBS leads. As the Neuropixels probe was inserted through the burr hole, there was only a limited number of areas and approaches using angles in three dimensions to insert the Neuropixels probe. **b**. Taking measures of the burr hole (upper right in **a**), we reconstructed the burr hole relative to the participant's brain (produced using FreeSurfer and MMVT[5–9], to identify the possible locations for the Neuropixels probe insertion taking into account the size of the 3D structure of the headstage and insertion into a cortical gyrus. Grey surface is the pial surface and white reconstructed volume is the white matter for the participant. **c-d**. The resultant putative location of the Neuropixels probe based on these measures in 3D (**c**) and overlaid on the MRI (**d**). **e-g**. Same process but for Pt. 02. **h**. Photographs of the open craniotomy (Pt. 03) and process of tracing the sulci to identify location of the craniotomy. **i**. Reconstructed brain with the overlaid trace of the craniotomy photograph of the lateral temporal lobe, with the putative location of the inserted Neuropixels probe shown on the brain. **j-k**. The resultant putative location of the Neuropixels probe based on the projected photograph to the 3D location on the brain.

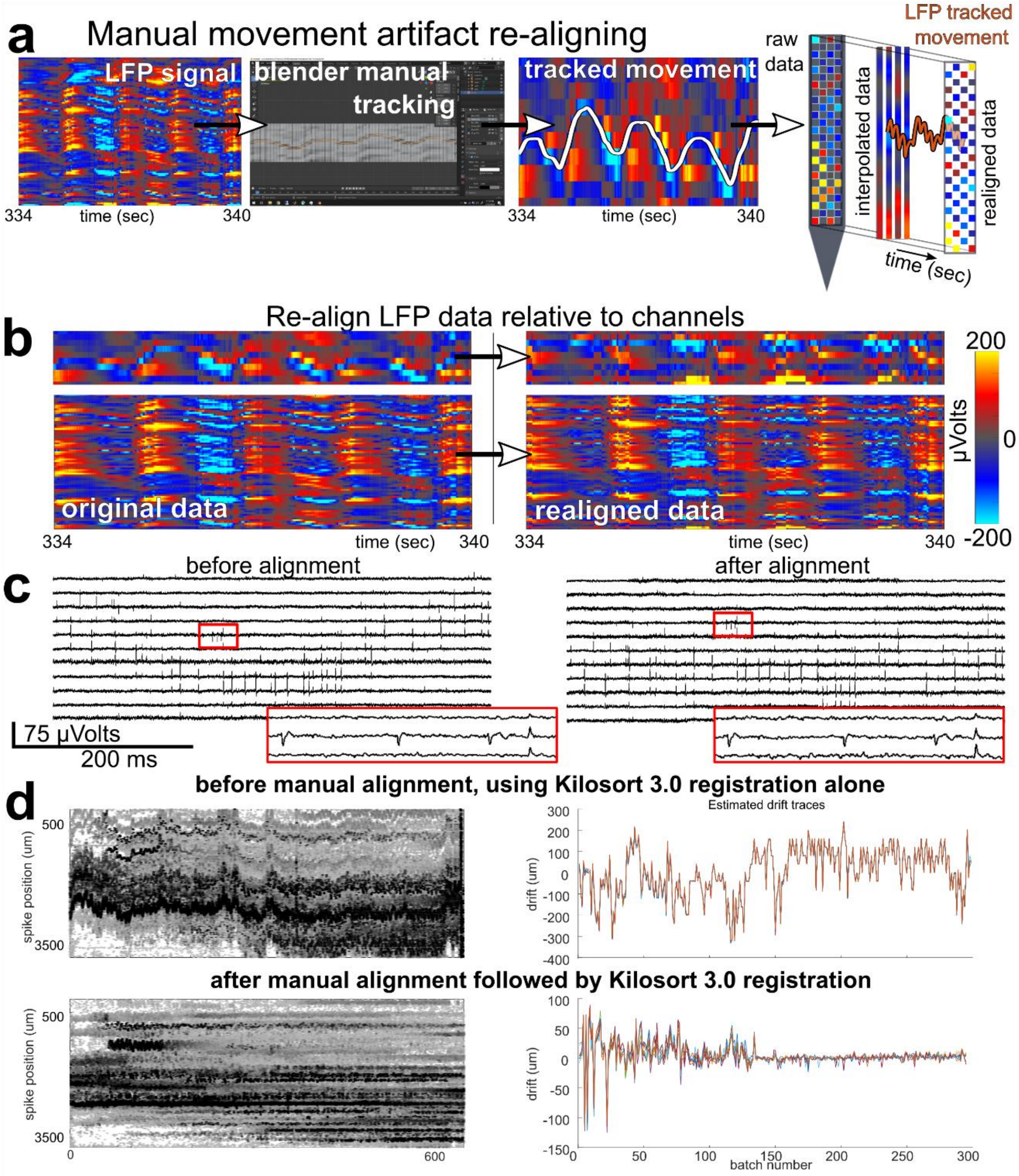


**Supplemental Figure 3: Realigning the data relative to heartbeat-induced movement artifact. a.** Illustration of evidence of tissue movement relative to the electrode recordings in the LFP (shown in red-blue color scale with the range in µvolts shown in **b**). This is quantified by manually tracing these “band shifts” using the Blender program, followed by detection of these movements in the LFP and tracking of these movements across channels (white line, second to rightmost plot). **b.** LFP before (left) and after (right) adjusting for movement effects. **c.** High frequency (action potential) frequency signal before (left) and after (right) adjusting for movement effects. **d**. Top row: Kilosort 3.0 registration and alignment alone could not compensate for the drift evident in the detected spike waveforms (left) and the estimated drift spanned hundreds of microns (right). Bottom row: manual alignment (**a-b**) followed by Kilosort 3.0 sorting resulted in improved spike alignment through time (left) and reduced drift (right).

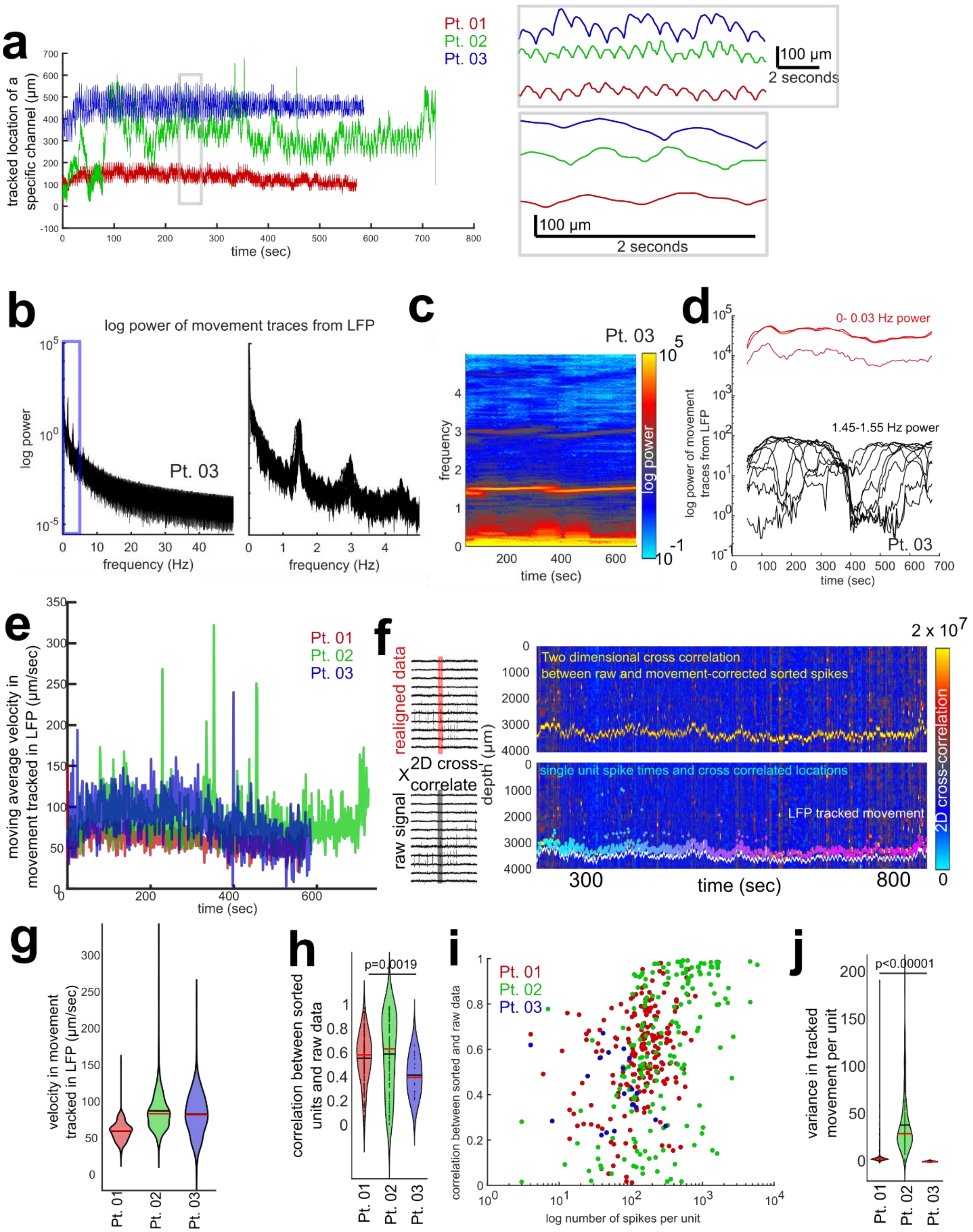

a
Pt. 01
Pt. 02
Pt. 03
tracked location of a specific channel (μm)
time (sec)
100 μm
2 seconds
b
log power of movement traces from LFP
log power
Pt. 03
frequency (Hz)
c
Pt. 03
frequency
log power
time (sec)
d
0- 0.03 Hz power
1.45-1.55 Hz power
log power of movement traces from LFP
Pt. 03
time (sec)
e
moving average velocity in movement tracked in LFP (μm/sec)
time (sec)
f
realigned data
X
2D cross-correlate
raw signal
depth (μm)
Two dimensional cross correlation between raw and movement-corrected sorted spikes
single unit spike times and cross correlated locations
LFP tracked movement
2D cross-correlation
2 x 10^7
time (sec)
g
velocity in movement tracked in LFP (μm/sec)
h
correlation between sorted units and raw data
p=0.0019
i
correlation between sorted and raw data
log number of spikes per unit
j
variance in tracked movement per unit
p<0.00001

**Supplemental Figure 4. Cross correlation between the interpolated sorted units and the raw data**. **a**. tracked movement from the LFP for the three participants, including zoomed in views in the grey boxes. **b**. Log power of the tracked movement from LFP for one participant for 0-50 Hz (power curves, left, and spectrogram, right). **c**. Log power of the tracked movement from LFP for one participant for 0-5 Hz (power curves, left, and spectrogram, right). **d**. Power of two different frequency bands (1.5 and 0.02 Hz) of the tracked movement from LFP through time for one participant (power curves). **e**. Moving average calculated velocity (µm/sec) of the movement as tracked in the LFP per participant. **f**. Left: We could correlate the 12 channels with the waveforms on a per-unit and per channel level and overlay the raw data (black lines) and the sorted movement tracked data (red lines). Right: Approach for cross correlating the movement-corrected neural data and the raw signals by cross correlating the 12 channels of the sorted spikes with the raw data with tracking of individual units along the raw channel data (top, middle, purple dots) based on the cross-correlation values (shown in color map) which was paired with the LFP tracked movement (white lines). **g**. Mean and median velocity (µm/sec) of the movement as tracked in the LFP per participant. **h**. Correlation values between sorted units and raw data per unit across 12 channels per unit (Kruskal-Wallis multiple comparisons test; Chi-sq.=12.29; p=0.0019). Scatter plots are individual unit correlation values per participant. **i**. Correlation values on the per-channel waveform basis between the interpolated and raw data sets versus the log spike rates per unit. Each dot indicates a unit. **j**. Mean variance in tracked cross-correlated single units. Asterisk indicates significant differences between each participant in the variance (Kruskal-Wallis multiple comparisons test; Chi-sq.=399.39; p<0.00001).

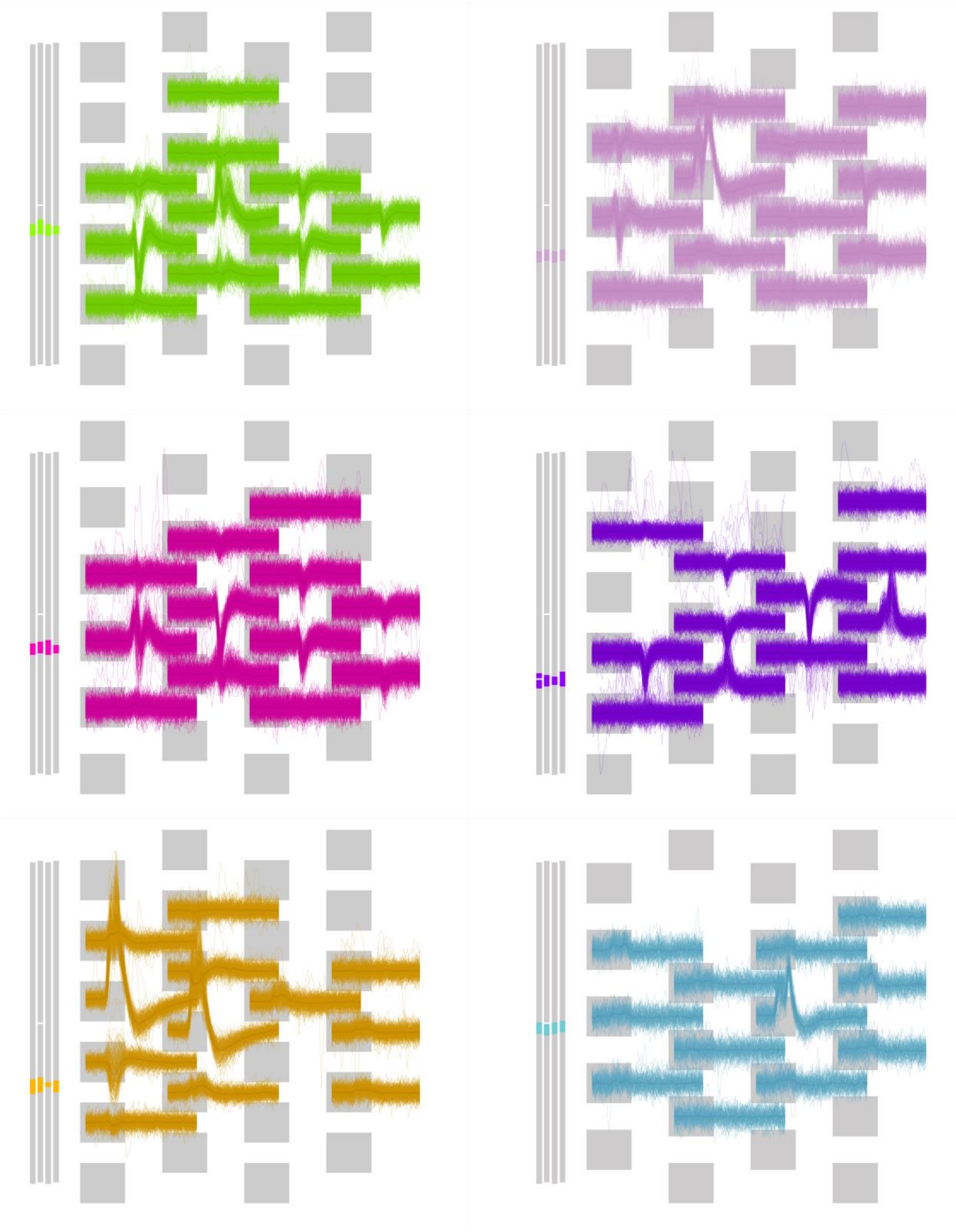

**Supplemental Figure 5. Example complex waveforms for six different units (each color-coded set of waveforms) across the data set**. Original waveforms are overlaid relative to the recorded channels, with the grey bars to the right indicating the location of the units along the Neuropixels probe.

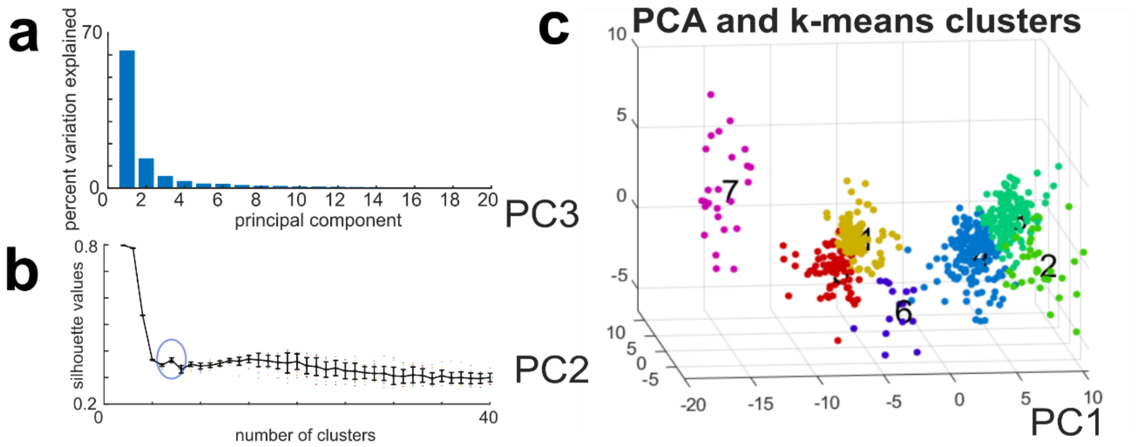


**Supplemental Figure 6. PCA with k-means clustering**. **a**. Percent variance explained by each principal component calculated across the first 6 channels per unit. **b**. Silhouette values for each k-means cluster number for the first 40 principal components after re-running the k-means clustering analyses 20 times (including 1000 iterations). **c**. Scatter plot of seven clusters (using k-means clustering) in the first three principal components. Color coding reflects different clusters.

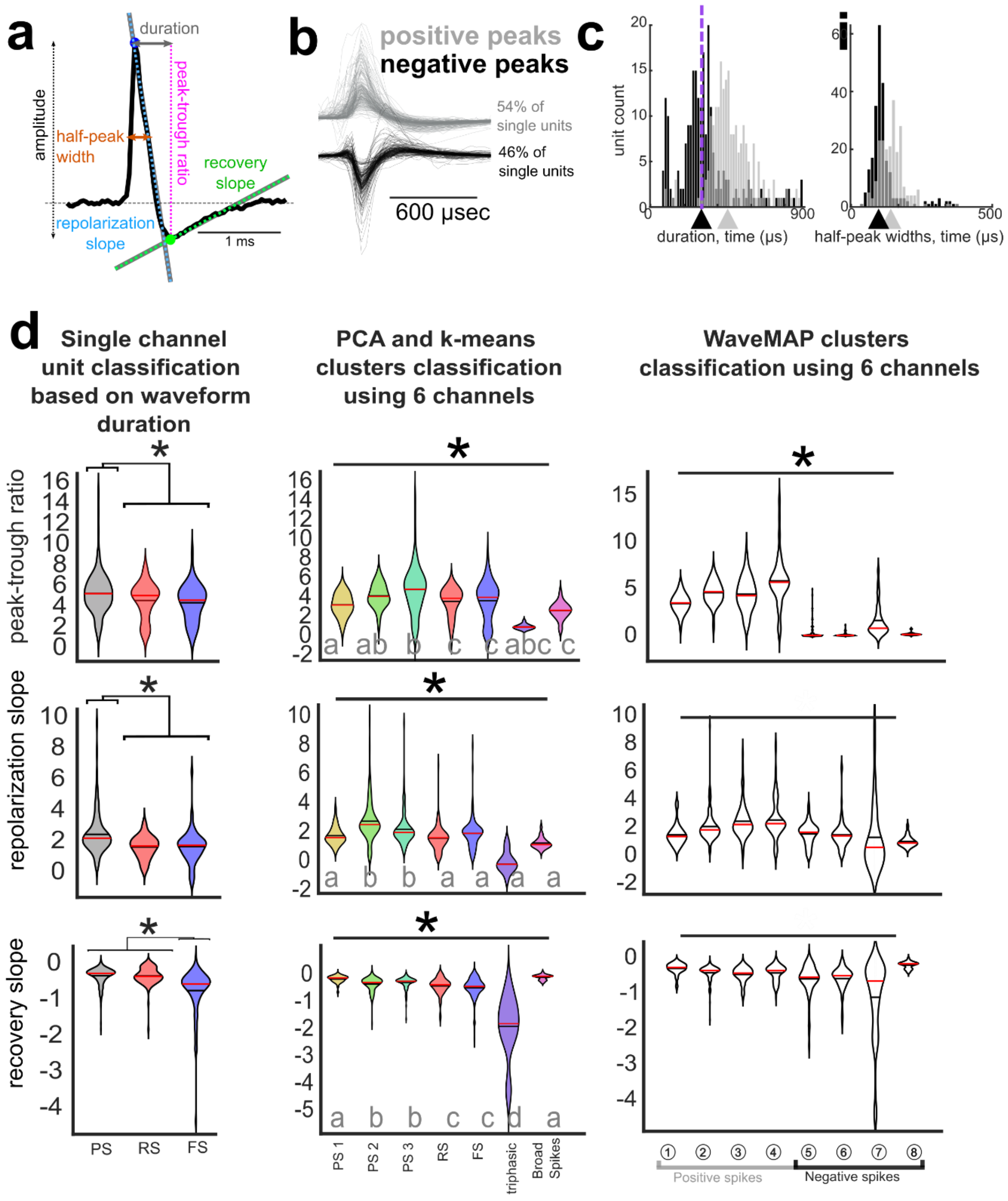


**Supplemental Figure 7. Waveform measures and PCA with k-means clustering**.

**a**. Waveform measures on an example, spike waveform including the spike duration, halfwidth, peak-trough ratio, repolarization slope, recovery slope, and amplitude measures. **b**. Largest waveforms per unit showing positive polarity (grey) and negative polarity (black) waveforms across the data sets. **c**. Distribution of positive polarity (grey) and negative polarity (black) half-peak width and spike duration (peak-trough duration) of the largest waveform per unit (Pt. 01-

03). **d**. The peak-trough ratio, repolarization slope, recovery slope of the channel with the largest waveform violin plots for the different clustered waveforms using the single channel and multichannel classifications. * indicate significant differences between all putative cell or waveform types, Kruskal-Wallis multiple comparison test, $p<0.001$ with *post hoc* Tukey-Kramer test. Grey letters a-c indicate statistically separable groups. Left: single channel classifications. Middle: multichannel classification based on the principal components and k-means clustering across patients. Right: multichannel classification based on the WaveMAP clustering across patients.

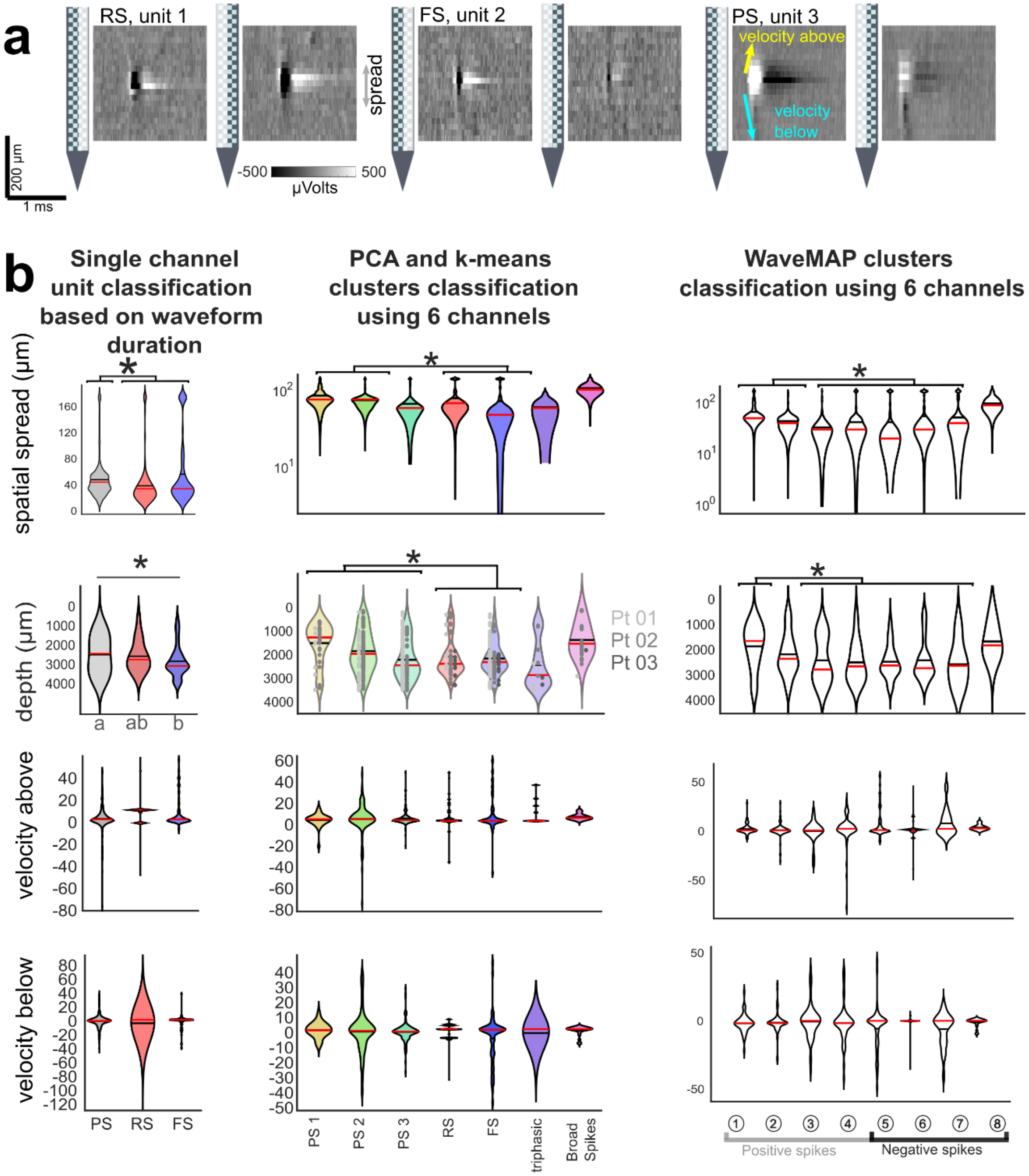


**Supplemental Figure 8. Waveform measures and PCA with k-means clustering**.

**a**. Spatial spread examples of individual units (averaged across all waveforms attributed to a given unit) across the left and right columns of the Neuropixels probe. Voltage indicated by the greyscale color scheme. Bottom plot: diagram (arrows) demonstrating the spatial mapping calculations, including spatial spread, velocity above (yellow arrow), and velocity below (cyan arrow) the center point (channel with the largest waveform). (Measures shown in **Supplemental Fig. 4**). **b**. The spatial spread, depth, velocity above, and velocity below the center point (channel with the largest waveform) violin plots are shown for the different waveform types. * indicate significant differences between all putative cell or waveform types, Kruskal-Wallis multiple comparison test, $p<0.001$ with *post hoc* Tukey-Kramer test. Grey letters a-c indicate statistically separable groups. Left: single channel classifications. Middle: multichannel classification based on the principal components and k-means clustering across patients. Right: multichannel classification based on the WaveMAP clustering across patients.

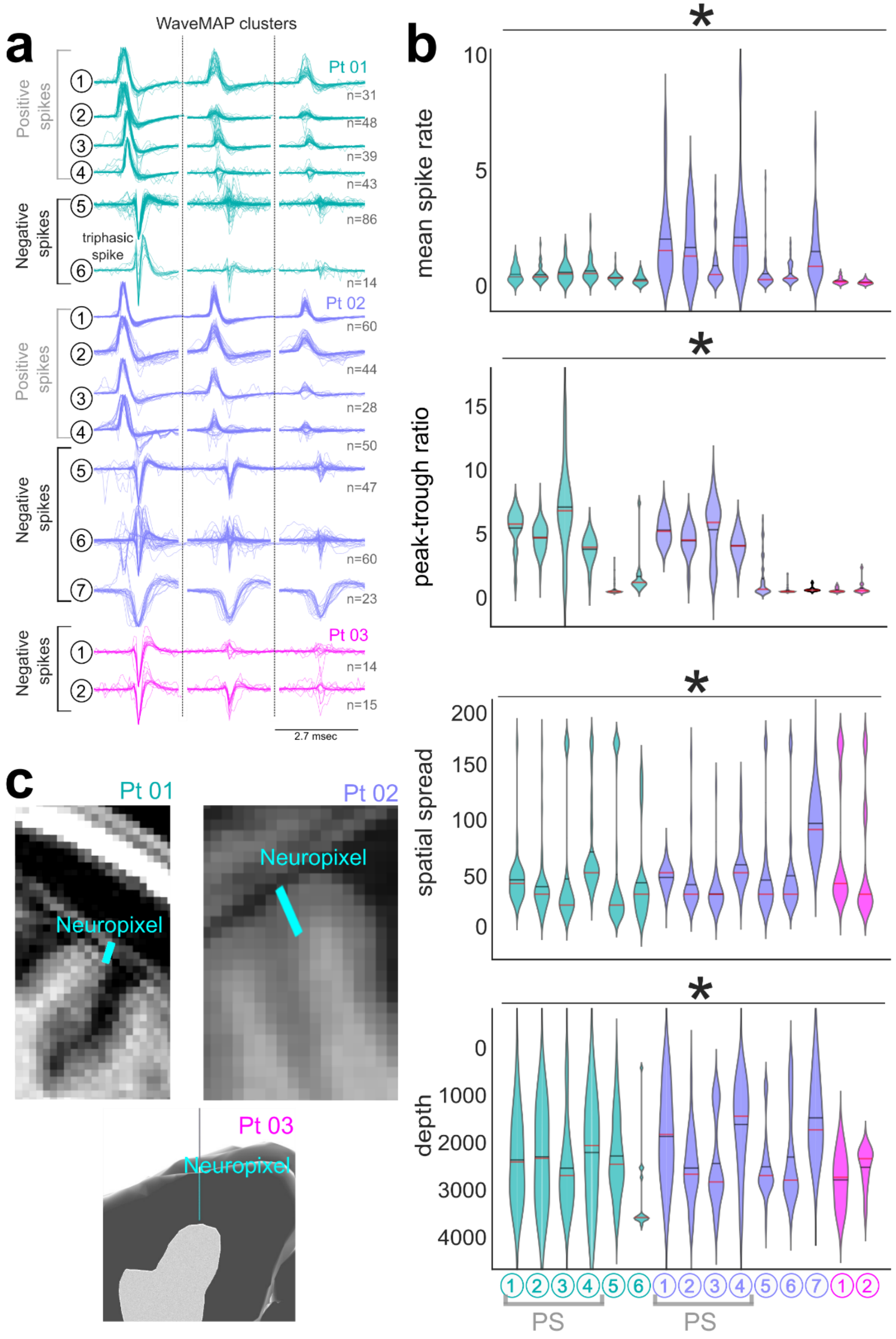


**Supplemental Figure 9. Waveform Features of Units Clustered with WaveMAP**.

**a**. Waveforms per cluster per patient (indicated as Pt. xx). The sorted clusters (with WaveMAP) was performed on the per-patient level which is in contrast with the WaveMAP clustering performed across all three participants (Fig. 3). **b**. The remaining measures are per patient, showing mean firing rate, peak-trough ratio, spatial spread, and depth violin plots for the different clusters. * indicate significant differences between putative cell or waveform types, Kruskal-Wallis multiple comparison test, $p<0.001$. **c**. Electrode locations relative to the cortical surface and cortical regions.

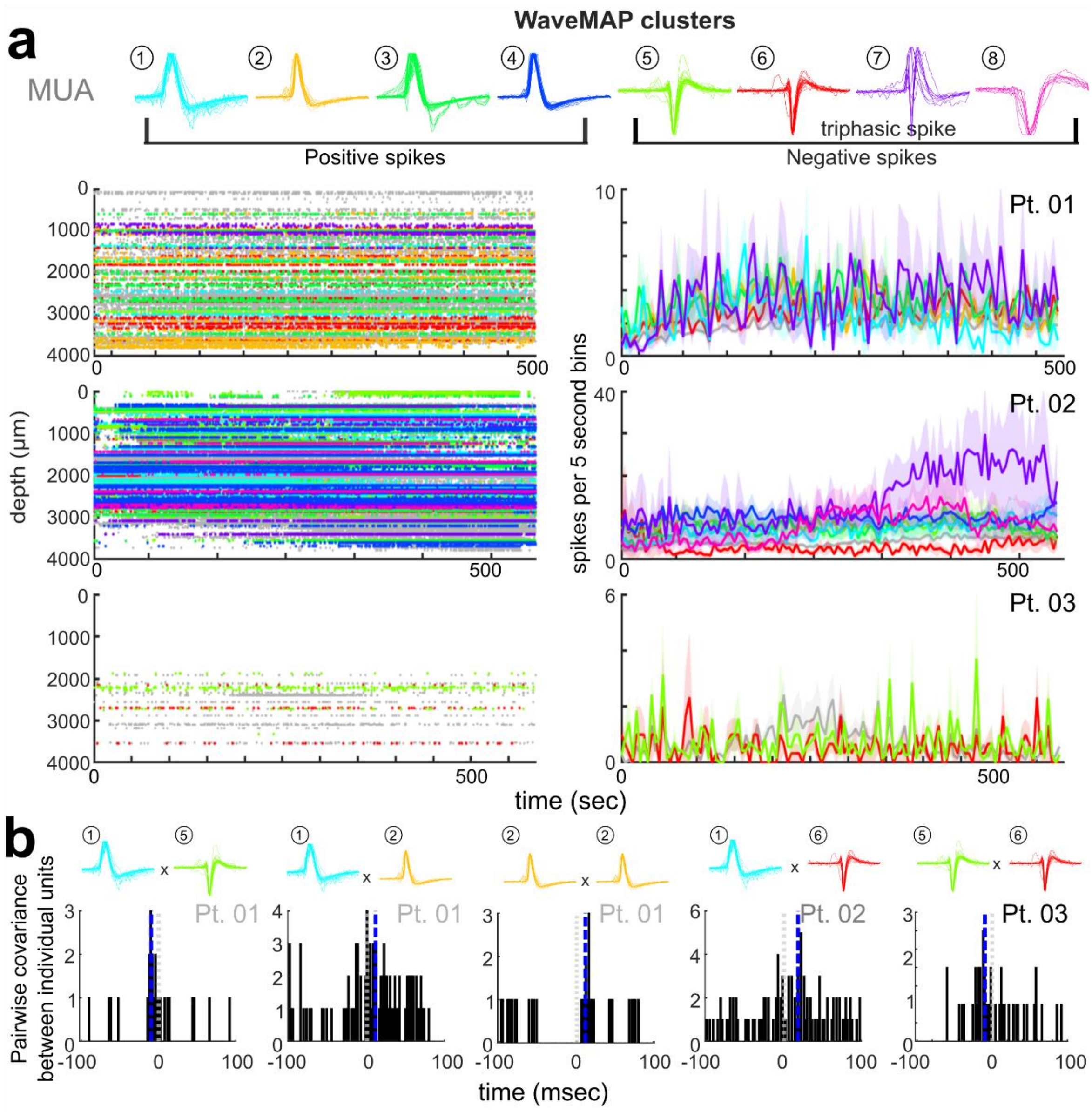


**Supplemental Figure 10. Single units through time and correlation relationships between units. a.** Left: the raster plots of spike times throughout the recording for the different classes of single units as clustered by WaveMAP (color coded relative to the waveform from the channel with the largest amplitude per unit) as well as the MUA activity (grey). Right: Spike counts binned in 5 second bins throughout the recordings for the three participants. **b**. Example cross-covariance of the spike times between individual units of different putative classes as clustered by WaveMAP (as titled). The vertical blue line shows the mean per plot.
